# Pervasive phenotypic effects of FBXO42 controlled by regulation of PP4 phosphatase

**DOI:** 10.1101/2025.05.02.651838

**Authors:** Hongbin Yang, Paul Smith, Yingying Ma, Emily Southworth, Varun Gopala Krishna, Beatrice Salerno, Joseph Rowland, Domenico Grieco, Iolanda Vendrell, Roman Fischer, Benedikt M Kessler, Vincenzo D’Angiolella

## Abstract

F-box proteins are the substrate recognition modules of the SCF (SKP1–Cullin– F-box) E3 ubiquitin ligase complex. FBXO42, an understudied member of this family, has recently emerged as a modulator of key cellular processes, including cell cycle progression, DNA damage response and glioma stem cell survival. In this study, we define the function of FBXO42 as a major regulator of the protein phosphatase PP4. Phosphoprotein phosphatases (PPPs) have a broad array of substrates, hence necessitating tight regulation. We observe that FBXO42 ubiquitinates the PP4 complex to govern the assembly of regulatory and catalytic subunits with the net effect of restraining the latter’s phosphatase activity. FBXO42 depletion unleashes PP4 activity with broad cellular effects highlighting FBXO42 as a novel regulatory node in ubiquitin-mediated signalling for future therapeutic exploitation.

**Impact Statement:** FBXO42 is a major regulator of PP4 phosphatase impacting multiple cellular phenotypes.

## Introduction

Modification by ubiquitin is a widespread signalling cascade to regulate essential biological processes (Damgaard, 2021). The system operates by the sequential addition of ubiquitin moieties to form chains on substrates which are recognized by the proteasome for degradation (Hershko & Ciechanover, 1992).

The fate of ubiquitinated proteins depend on a code largely reliant on polyubiquitin chain topologies thus generated (Komander & Rape, 2012). Ubiquitin moieties linked through lysines 48 or 11 are typically degradative signals, while polyubiquitin chains harbouring alternate ‘atypical’ linkages have other distinct outcomes. In numerous instances, non-canonical polyubiquitin chains serve as scaffolds to recruit protein complexes (Agrata & Komander, 2025).

The specificity in the ubiquitin system is dictated by a hierarchical system of three components: ubiquitin activating (E1), conjugating E2 and ligase (E3s) enzymes. E3s are comprised of large families of enzymes which select substrates based on specific assembly modules (Morreale & Walden, 2016). Most abundant among the E3s, Cullin Ring Ligases (CRLs) use a central component (cullin) to bridge a substrate recruitment protein and a RING Finger containing protein which, in turn, recruits E2 enzymes for ubiquitin transfer (Harper & Schulman, 2021; Zheng et al., 2002). F-box proteins are the substrate adaptor modules of cullin 1 (Cul1) complexes, the module being composed of S-phase kinase-associated protein 1 (Skp1), Cul1 and the F-box itself (SCFs). These are further subclassified as F-boxes with a WD-40 domain (FBXWs), F-boxes with a leucine zipper domain (FBXLs), F-boxes with other domain (FBXOs) based on their substrate recruitment domains (Cardozo & Pagano, 2004; Fouad et al., 2019).

Prototypical F-box proteins engage substrates by binding to discrete sequences called “degrons”. As an example, Cyclin F (FBXO1) recognizes a bivalent domain (F-deg) composed of a phosphorylated residue and a cyclin binding domain (RxIF) (Ngoi et al., 2025; Yang et al., 2024). Most of the substrates are recognised in G2/M when the level of cyclin F itself peaks, allowing the control of multiple substrates by a single E3. As exemplified above, there is significant interplay between phosphorylation and degradation with some F-boxes (particularly Fbxws) recognising substrates only after phosphorylation.

Phosphorylation is a reversible modification balanced by the opposing action of kinases and phosphatases. PPPs (PP1-PP7) use a common catalytic process which entails activation of a water molecule acting as the nucleophile in the dephosphorylation reaction (Shi, 2009). Albeit similar in mechanism of action, PPPs differ significantly in subunit composition and assembly. PP4 uses a total of five regulatory subunits to assemble distinct hetero-oligomeric complexes. Indeed, PP4C (the catalytic subunit) can assemble with PP4R1 or PP4R2, PP4R3A, PP4R3B or PP4R4 thus leading to the formation of four distinct oligomeric complexes (PP4C/PP4R1; PP4C/PP4R2/PP4R3A; PP4C/PP4R2/PP4R3B; PP4c/PP4R4) (Chen et al., 2008; Hwang et al., 2016; S. P.

Lyons et al., 2021). Elegant structural studies have defined the mode of substrate recognition operated by PP4R2 in complex with PP4R3A, recognising proline rich residues in targeted substrates (Ueki et al., 2019). The oligomeric composition and diversity of PPPs are required for substrate selectivity, but little is known about the substrate engagement strategy and the mechanisms to regulate oligomeric assembly of PP4.

We identify the E3 ligase adaptor FBXO42 as a major regulator of the phosphatase PP4. Our study highlights that SCF^FBXO42^ induces non-degradative ubiquitination of PP4, regulating PP4R2 subunit assembly with PP4C, thus, acting as a major inhibitor of PP4 phosphatase activity. Depletion of FBXO42 results in widespread effects on phosphorylation brought about by spurious activation of PP4. This mechanism highlights a yet undiscovered signalling node with multiple effects on cell morphology, survival and proliferation.

## Results

### Loss of FBXO42 induces multiple cellular defects including defective DNA damage response (DDR)

The advent of Clustered Regularly Interspaced Short Palindromic Repeats (CRISPR) has broadly enhanced our capacity for querying drug-gene interactions at a genome-wide scale. Several screens have been performed to identify genes contributing to cell viability upon the inhibition of the Ataxia telangiectasia and Rad3-related protein kinase (ATR) (Saldivar et al., 2017; Saldivar et al., 2018), a key controller of the DNA damage response (DDR) and regulator of mitotic entry at the G2/M checkpoint. A consensus set of vulnerabilities became evident from four genome-wide CRISPR screens using two ATR inhibitors (AZD6738 and VE-821) in multiple cell lines (Hustedt et al., 2019). FBXO42 was identified among the significant genes whose depletion increased cell death upon ATR inhibition using two inhibitors in the four indicated cell lines (Figure 1A and Figure S1A) (Hustedt et al., 2019; Wang et al., 2019). A screen focused on the ubiquitin system testing the genes mediating sensitivity to 41 compounds, identified FBXO42 as a vulnerability in cells treated with microtubule depolymerizing agents (colchicine) and BI-2536, a specific inhibitor of the mitotic kinase Polo-like kinase 1 (Plk1, Figure 1B and S1B) (Hundley et al., 2021). Furthermore, the loss of FBXO42 was shown to promote resistance to MEK inhibitors in melanoma (Nagler et al., 2020) and specifically impair the growth of a subset of glioblastoma stem-like cells (Hoellerbauer et al., 2024). We had previously conducted a high resolution CRISPR screen to identify genes controlling cell viability in response to treatment with ionizing radiation (IR) (Yang et al., 2024), where FBXO42 was also among the hits noted (Figure 1C).

**Fig. 1.**
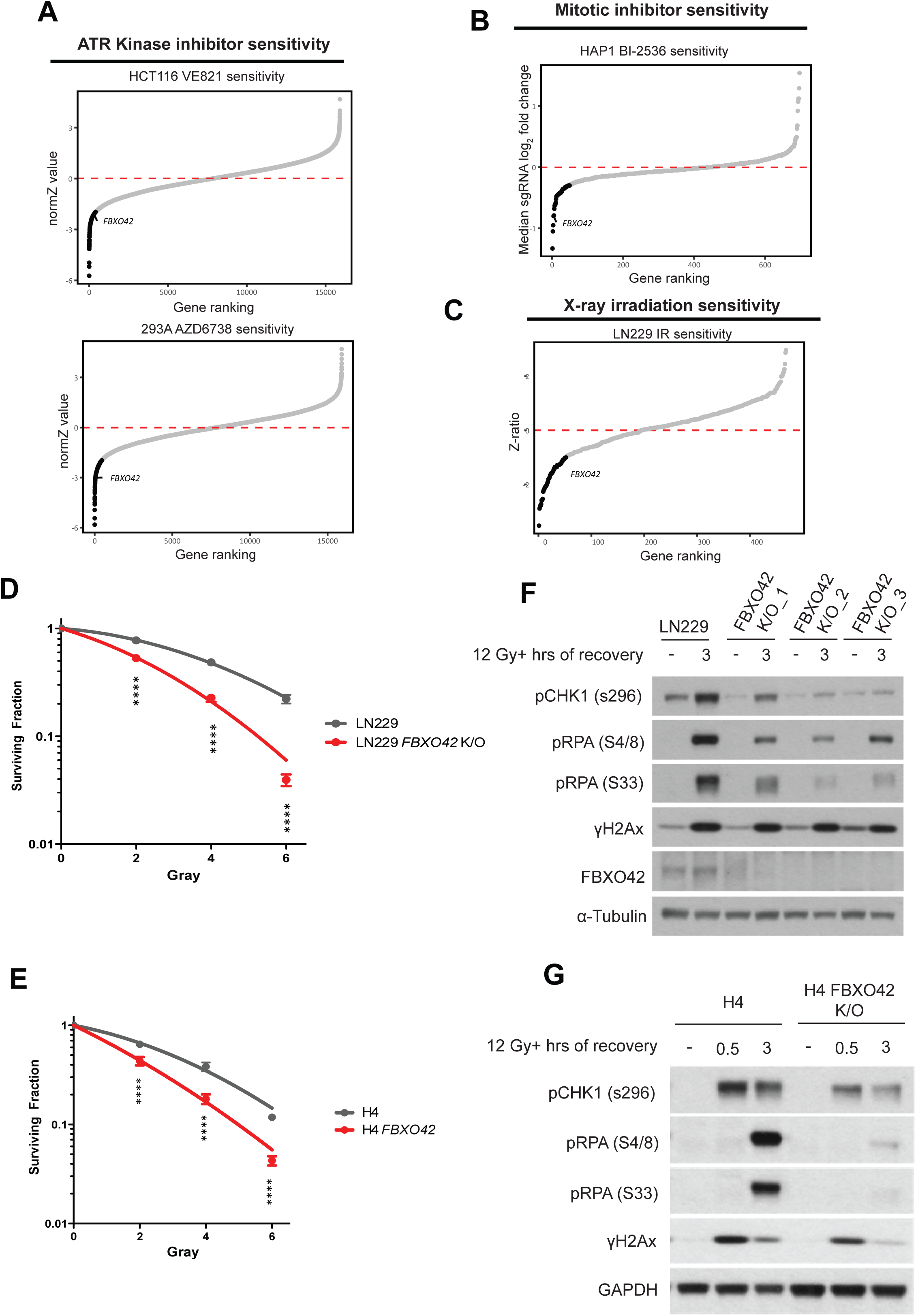
Loss of FBXO42 induces multiple cellular defects including defective DDR responses. A. CRISPR screen probing vulnerabilities in response to inhibitors of the ATR kinase from (Hustedt et al., 2019; Wang et al., 2019) in HCT116 and 293A cells. Outcome of screens are represented by gene ranking using Z-Score (normZ) values, using a significance cut-off of ≤ -1.96. FBXO42 is highlighted. B. CRISPR screen probing vulnerabilities in response to inhibitor of the kinase PLK1 from (Hundley et al., 2021) in HAP1 cells. Outcome of screen is represented by gene ranking using median sgRNA log2 fold change cut-off of ≤ -0.3, FBXO42 is highlighted. C. CRISPR screen probing vulnerabilities in response to IR (Yang et al.,2024) in LN229 cells. Outcome of screen is represented by gene ranking using Z-ratio values, using a significance cut-off of ≤ -1, to reflect the use of a focussed CRISPR-Cas9 screen in place of genome-wide. FBXO42 is highlighted. D. Colony formation assay in FBXO42 K/O cells. LN229 parental cells or LN229 FBXO42 K/O cells were seeded for colony formation assay and challenged with the indicated dose of IR. Seven days after IR, cells were stained with crystal violet and counted. Error bars represent SDs of three biological replicates. Two-tailed unpaired *t* test was performed as statistical analysis. **P ≤ 0.01; ***P ≤ 0.001; ****P ≤ 0.0001. E. Immunoblotting after IR treatment and 3 hour recovery in LN229 parental and FBXO42 K/O cells. DNA damage markers detected: pCHK1 S296 (cell cycle checkpoint), pRPA32 S4/8 (ssDNA at DSB), pRPA32 S33 (single-strand breaks), and γH2Ax (DNA DSBs). F. Same as D in H4 parental and H4 FBXO42 K/O cells. G. Same as E in H4 parental and H4 FBXO42 K/O cells.

To exclude that FBXO42 is not a frequent hitter due to an inherent characteristic of the sgRNA targeting it, we generated two glioblastoma cell lines (LN229 and H4) where FBXO42 was knocked out using sgRNA sequences distinct from the library (Appendix S1). Both LN229 and H4 cell lines lacking FBXO42 were significantly more sensitive to VE-821 (ATRi) and indibulin (a microtubule depolymerising agent analogue of colchicine), confirming the functional role of FBXO42 after ATR inhibition and mitotic spindle checkpoint activation identified in the genome-wide screens (Figure S1C, S1D, S1E and S1F). Additionally, sensitivity to IR was also evident through classical clonogenic assays in both LN229 and H4 cells lacking FBXO42 (Figure 1D and E).

Upon analysing the phosphorylation events induced by IR treatment in cells lacking FBXO42, we observed a reduction in phosphorylation events controlling the DNA damage checkpoint (levels of pChk1 S345) after IR treatment. Also, we noted a decrease in RPA32 phosphorylation at serines 4/8 [marker of ssDNA at DSB repair sites phosphorylated by Ataxia telangiectasia mutated (ATM) and DNA-dependent protein kinase (DNA-PK)] and serine 33 (marker of ssDNA phosphorylated by ATR) (Figures 1F and 1G). γH2Ax levels induced by IR were not impacted by depletion of FBXO42 (Figures 1F and 1G). The overall reduction of phosphorylation events is indicative of a defect in DNA damage signalling rather than a compromised DNA repair.

### FBXO42 interacts with PP4 phosphatase

The multitude of phenotypic effects induced by FBXO42 depletion could be explained by the fact that often E3 ligases target multiple substrates by recognizing degrons in proteins irrespective of the biological process they are involved in. A well-established example is beta-transducin repeat containing E3 ubiquitin protein ligase (β-TrCP) which recognizes the DSGXXS degron once both serines have been phosphorylated (Busino et al., 2003; Guardavaccaro et al., 2003; Jin et al., 2003; Margottin-Goguet et al., 2003). The control of degradation is dictated by kinases which phosphorylate the DSGXXS, with β-TrCP controlling multiple substrates involved in diverse cellular processes (Bi et al., 2021). We reasoned FBXO42 might follow a similar principle and attempted to identify interactors of FBXO42 which are specifically impacted by IR, given its functional role in controlling the DDR. To isolate FBXO42 substrates we treated cells with MLN4924 (a selective inhibitor of NAE1) (Soucy et al., 2009) and identified interactors after immunoprecipitation and liquid chromatography/mass spectrometry (LC/MS) as previously done for other CRLs (Burdova et al., 2019; Chen et al., 2022; Fung et al., 2018; Raducu et al., 2016). Neddylation is required for ubiquitination mediated by CRLs (Duda et al., 2008), thus inhibition of NAE1 results in inactivation of CRL’s ubiquitination activity. Differential interacting partners enriched in the presence of MLN4924 are likely substrates, as their interaction with the adaptors is increased when the CRL mediated ubiquitination is prevented. Surprisingly, IR treatment alone does not lead to major differences in interactions with Fbxo42 (Figure 2A, *lower right quadrant*). As expected, MLN4924 reduces the interaction of SCF^FBXO42^ with Nedd8, judging by its enrichment in the top right quadrant (Figure 2A). Most importantly, in the presence of MLN4924 FBXO42 binds more strongly to the protein phosphatase 4 (PP4) complex, including subunits PP4C, PP4R1, PP4R2, PP4R3A (SMEK1) and PP4R3B (SMEK2) together with CCDC6 (Cerrato et al., 2018) (Figure 2A).

**Figure 2.**
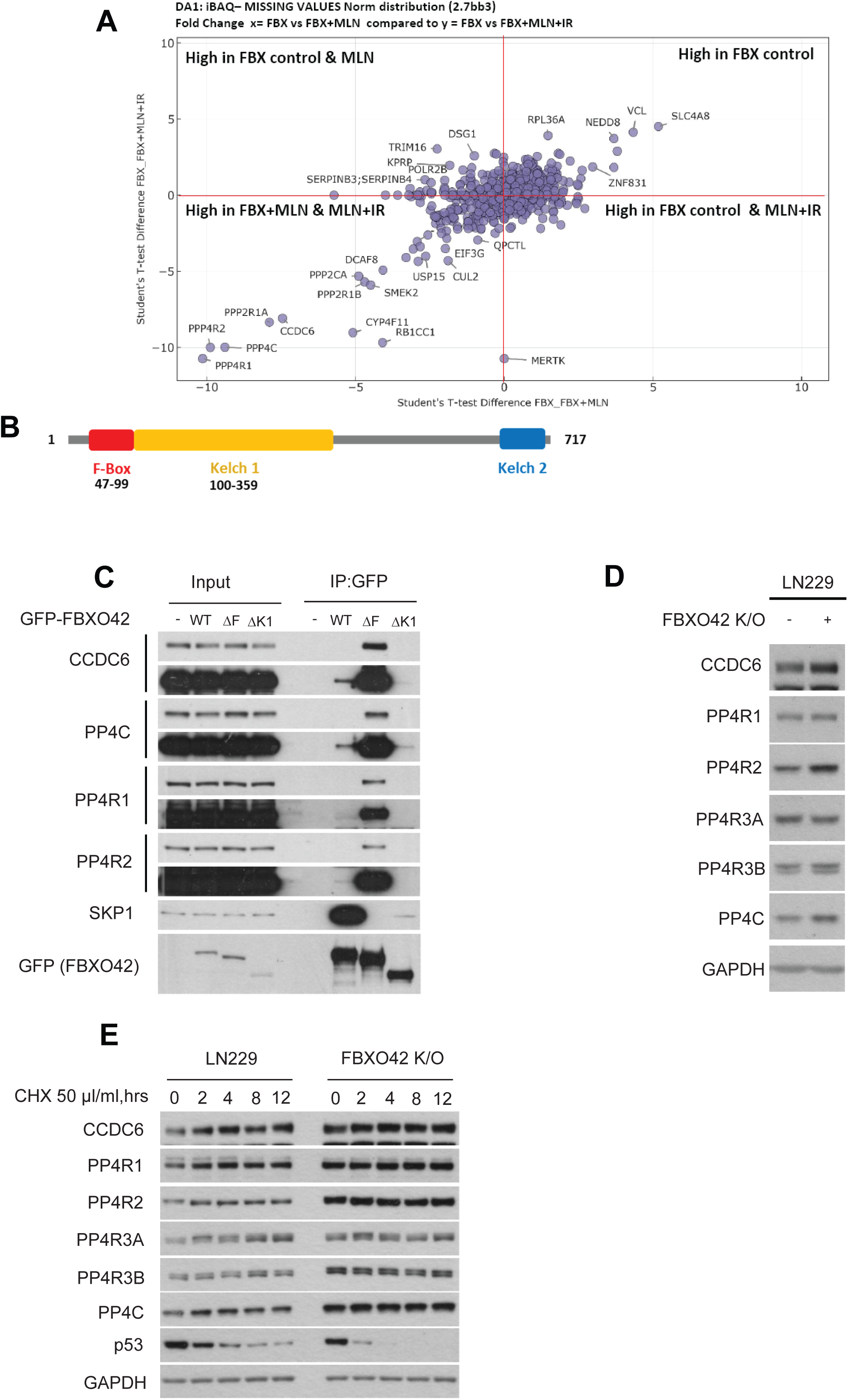
FBXO42 interacts with PP4 phosphatase. A. Scatter plot of the log fold changes of FBXO42 IP-MS experiments +/- MLN4924 vs +/- ionisation radiation. The condition and relative level of the proteins is indicated in each quadrant. B. Schematic representation of the domains in FBXO42. The F-box domain and Kelch domain are highlighted. The kelch domain is divided in two sections (Kelch1 and Kelch2) as reported on PFAM. C. Immunoblotting after immunoprecipitation of GFP FBXO42 WT, F-box (DF) mutant and Kelch (DK1) mutant. Input samples are presented on the left. D. Immunoblotting of the indicated proteins in LN229 parental and FBXO42 K/O cells. E. Immunoblotting of FBXO42 interactors after treating LN229 parental or Fbxo42 K/O cells with cycloheximide (CHX) 50ug/ml for the indicated times (hrs = hours).

In a parallel approach we established a cell line expressing FBXO42 fused to TurboID (an engineered biotin ligase that conjugates biotin to proteins) to identify proteins in proximity of FBXO42 (Figure S2A). This approach, which exploits a nonspecific biotin ligase, has been used extensively to identify interacting partners of cellular proteins (May & Roux, 2019) with improvements made to enhance efficiency (Larochelle et al., 2019). Also, in this approach we retrieved the PP4 phosphatase complex as an MLN4924 dependent interacting partner of FBXO42 (Figure S2A).

Like other F-box proteins FBXO42 has an F-box domain at the N-Terminus and a substrate recognition domain (Kelch domain) at the C-terminus (Figure 2B) (Cardozo & Pagano, 2004). To establish the domains involved in the recruitment of PP4 complex, we performed co-immunoprecipitation of either WT FBXO42 or mutants lacking either the F-box (ΔF) or the Kelch 1 (ΔK1) domain. As expected, ΔF mutants although unable to bind Skp1 and Cul1, strongly bound to the PP4 phosphatase, thus suggesting that PP4 <u>is</u> a *bona fide* substrate of FBXO42 (Figure 2C). The ΔK1 truncation, however, abolished binding to PP4, confirming the need for kelch domain in substrate engagement (Figure 2C).

We initially hypothesized that FBXO42 targeted PP4 phosphatase for ubiquitin dependent degradation and measured the levels of PP4 phosphatase subunits in cells where FBXO42 had been knocked out by CRISPR. We noted a consistent increase in PP4C and PP4R2 levels upon FBXO42 depletion, while other subunits were not impacted (Figure 2D). Simultaneously and surprisingly, no change in half-lives were noted for PP4 subunits in FBXO42 K/O cells (Figure 2E). In the same settings we also observed a reduction in the half-life of p53, which is mutated in LN229 cells. This observation has previously been reported, confirming the conditions of the experiment and common effects due to FBXO42 loss (Hoellerbauer et al., 2024; Lu et al., 2024). This phenomenon could be due to a cell line specific effect; thus, we measured PP4 half-lives in H4 cells harbouring a WT p53. In this case, no change in half-lives were observed for p53 or PP4 subunits (Figure S2B). Finally, overexpression of FBXO42 (in HEK293T cells) did not lead to significant changes in the levels of PP4 phosphatase (Figure S2C).

### FBXO42 ubiquitinates PP4 phosphatase

Based on its enrichment upon MLN4924 treatment along with unchanged half-lives in the absence of FBXO42, we predict PP4 complex to be ubiquitinated by Fbxo42 although not targeted for degradation. To further explore the latter, we tested the ability of FBXO42 to ubiquitinate PP4 directly. Expression of FBXO42 induced a significant increase in ubiquitination of all components of the PP4R2 complex including PP4C, PP4R2, PP4R3A and B (Figures 3A, S3A). To confirm that the detected ubiquitin ladder is dependent on FBXO42 activity, we tested ubiquitination of PP4 using FBXO42 mutants mentioned above (ΔF and ΔK1; Figure 2C). Both mutants were unable to ubiquitinate PP4 as the FBXO42 WT did (Figure S3B).

**Figure 3.**
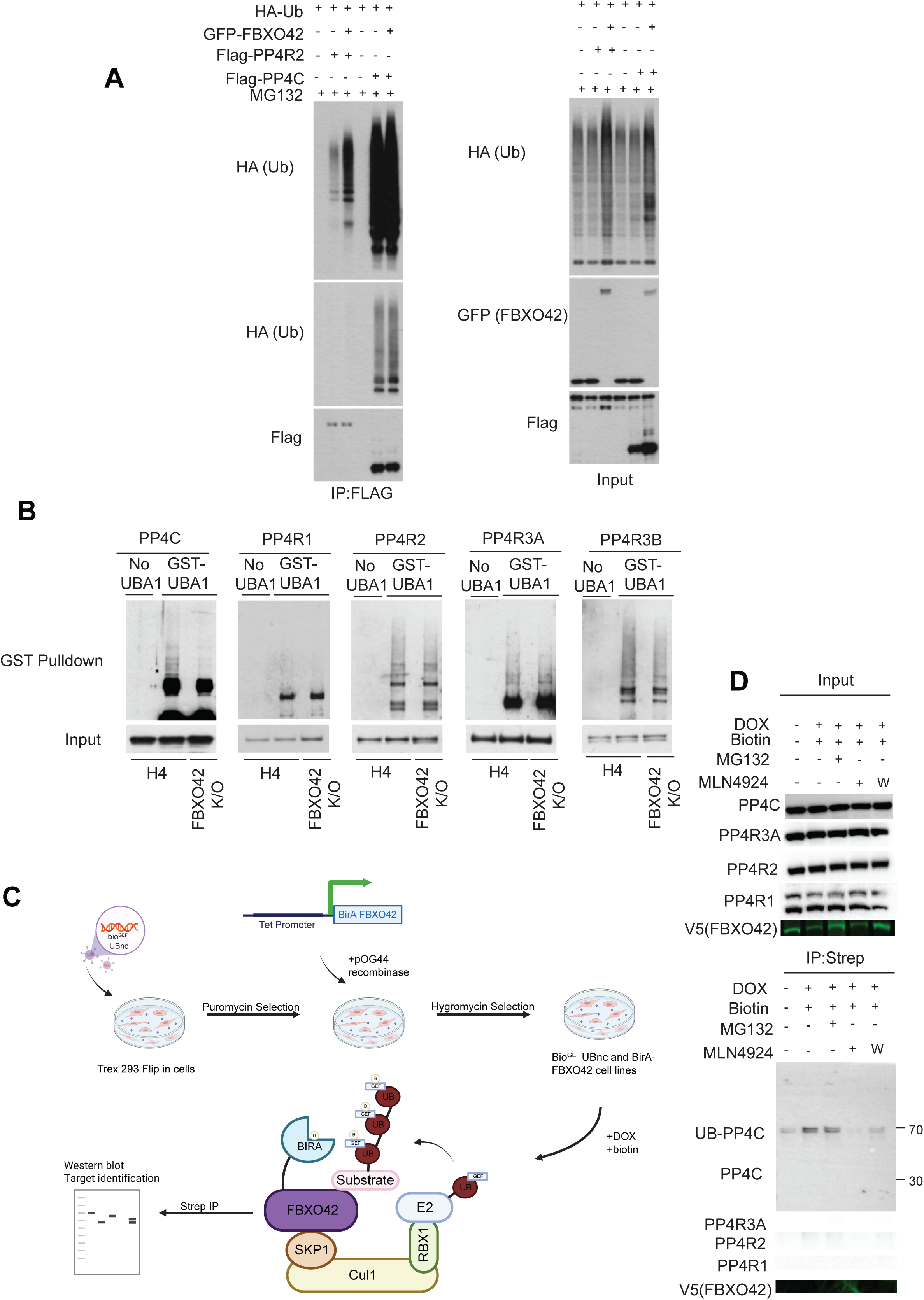
FBXO42 ubiquitinates the PP4 phosphatase. A. Immunoblotting after expression of GFP-FBXO42, HA-ubiquitin and either Flag-PP4R2 or Flag-PP4C in HEK293T as indicated. Flag-PP4R2 or Flag-PP4C was isolated *via* Flag agarose beads pulldown before immunoblotting. Input samples are in the right panel. B. Immunoblotting after isolation of endogenous ubiquitinated proteins using recombinant GST-tagged UBA domain of UBIQLN protein from H4 parental cells or cells FBXO42 K/O. Input samples before immunoprecipitation are indicated – *bottom panel*. C. Schematic representation of the E-Stub proximity ubiquitin labelling and enrichment. BirA-FBXO42 was co-expressed with a biotin acceptor ubiquitin mutant (bio^GEF^UBnc) followed by biotin treatment. Nascent ubiquitinated substrates are thus biotinylated and immunoprecipitated. D. Immunoblotting pf PP4 complex after expression of (doxycycline-inducible) BirA– FBXO42 and biotin bio^GEF^UBnc-Ub in T-REx™ Hek293 cells post. Proteins were induced by doxycycline for 24 hours and supplemented with biotin for 3 hours with or without MG132/MLN4924 as indicated, before protein lysis and immunoprecipitation. Input samples indicated in top panel.

The non-physiological expression of E3 ligases and ubiquitin itself can lead to artefactual ubiquitination events. Especially in the case of protein complexes, overexpression might force ubiquitination of components of the complex that normally are not modified. We therefore used two parallel approaches to establish the direct ubiquitination targets: we first sought to detect PP4 ubiquitination at endogenous levels in WT and FBXO42 K/O cell lines and follow this up with a recently developed proximity ubiquitination assay called E3-substrate tagging by ubiquitin biotinylation (E-Stub) (Barroso-Gomila et al., 2023; Huang et al., 2024; Laura Merino-Cacho et al., 2025). Upon unbiasedly enriching all ubiquitinated proteins from parental LN229 cells using the recombinant ubiquitin-associated domain (UBA domain) of the UBQLN1 protein in a ubiquitin binding entity (UBE) pulldown assay (Fiil et al., 2013), we detected endogenous polyubiquitinated PP4 in LN229 control cells and compared it to FBXO42 K/O (Figure 3B). The same experiment was also conducted in H4 cell lines (Figure S3C). Taking together the results from LN229 and H4 on the different PP4 subunits in parental versus FBXO42 K/O lines we could conclude that: PP4R1 was unlikely to be ubiquitinated; the PP4 complex most impacted by FBXO42 depletion was the one containing PP4C-PP4R2-PP4R3A; and finally, the most consistent effect across the two cell lines was on the PP4C subunit (Figure 3B and S3C). Thus, we hypothesize that FBXO42 specifically ubiquitinates the PP4C subunit within the PP4R2 complex.

More evidence in support of the latter was revealed by our modified E-stub. In our experiment, we used an ubiquitin moiety with a biotin acceptor tag from (Barroso-Gomila et al., 2023; L. Merino-Cacho et al., 2025) and the E3 (FBXO42) with the biotin donor (BirA) (May & Roux, 2019). Both the acceptor and donor biotin module are under the control of tetracycline in a Flip-in system line (Figure 3C). Use of E-stub confirmed that we could retrieve ubiquitinated PP4C in proximity of the FBXO42 complex but not PP4R1 and PP4R2 (Figure 3D). The proximal ubiquitination of PP4C was dependent on the activity of FBXO42 as MLN4924 abolished the retrieval of PP4C. In addition, MG132 did not increase PP4C retrieval, a further proof that ubiquitination of PP4C did not lead to degradation. Interestingly, the detected PP4C band that was enriched by biotin had a higher molecular weight than what was visible in the input, suggesting an attachment of perhaps three ubiquitin moieties thus indicating a specific signalling event rather than a degradative signal (Figure 3D). In summary, the main interactor and substrate of FBXO42 is the PP4 phosphatase, suggesting the possibility that multiple phenotypic effects attributed to FBXO42 might be due to its modulation of this versatile phosphatase.

### FBXO42 is a major regulator of PP4 phosphatase

We have thus far observed that FBXO42 ubiquitinates PP4C and is, likely to regulate the activity of PP4 complex rather than the levels. For a comprehensive understanding of the effect of FBXO42 on phosphorylation, we performed a mass spectrometry based phospho-proteomic analysis in LN229 cells (parental and FBXO42 K/O).

The analysis of total protein levels was conducted <u>alongside</u> phospho-proteomic analysis to normalise effects induced by protein level alterations. FBXO42 K/O cells had a significant dysregulation of proteins (∼3000 proteins were either upregulated or downregulated) likely an indirect consequence of a broader dysregulation of phosphorylation. The levels of PP4 phosphatase components were not majorly impacted (Figure S4A), except for a slight upregulation of PP4C and PP4R2, also noted before (Figure 2D). A gene enrichment analysis was conducted to highlight the pathways being downregulated in FBXO42 K/O using EnrichR and selecting the MSigDB hallmarks 2020 enrichment tool (Figure S4B). This analysis highlighted significant alterations of genes involved in G2/M checkpoint, mitotic spindle and DNA repair in line with the phenotypes described above and previous publications (Hoellerbauer et al., 2024; Hundley et al., 2021; Nagler et al., 2020; Toledo et al., 2015). The upregulated pathways include oxidative phosphorylation and epithelial-mesenchymal transition (Figure S4C), important cancer associated phenomena which require further investigation. It is plausible that the observed alterations in protein levels are an indirect consequence of altered regulation of phosphorylation events.

The phospho-proteomic analysis revealed a significant difference between parental and FBXO42 K/O cells with 89 proteins showing increased phosphorylation and 45 proteins showing decreased phosphorylation (Figure S4D). Gene enrichment analysis of proteins whose phosphorylation was reduced revealed mitotic progression and the G2/M checkpoint (Figure S4E), in line with previously observed phenotypes ascribed to loss of FBXO42. The decrease in phosphorylation could be attributed to spurious activation of PP4 phosphatase leading to increased dephosphorylation. However, a challenge we encountered in this regard is a dearth in our understanding of specific substrates of PP4; comprehensive studies on a total proteome level showing changes in phosphorylation, in cells devoid of PP4 activity are lacking and only one study thus far has highlighted proline rich sites (FXXP and MXXP) as recognition substrates for the PP4R2 complex.

To reconcile this, we matched the identified PP4R2 substrates in (Ueki et al., 2019) with proteins whose phosphorylation decreased upon FBXO42 K/O. We observed that out of the 14 substrates of PP4R2 reported, 10 were upregulated in FBXO42 K/O (Figure 4A). In other words, about 71% of the known PP4R2 substrates are dephosphorylated in FBXO42 K/O.

**Figure 4.**
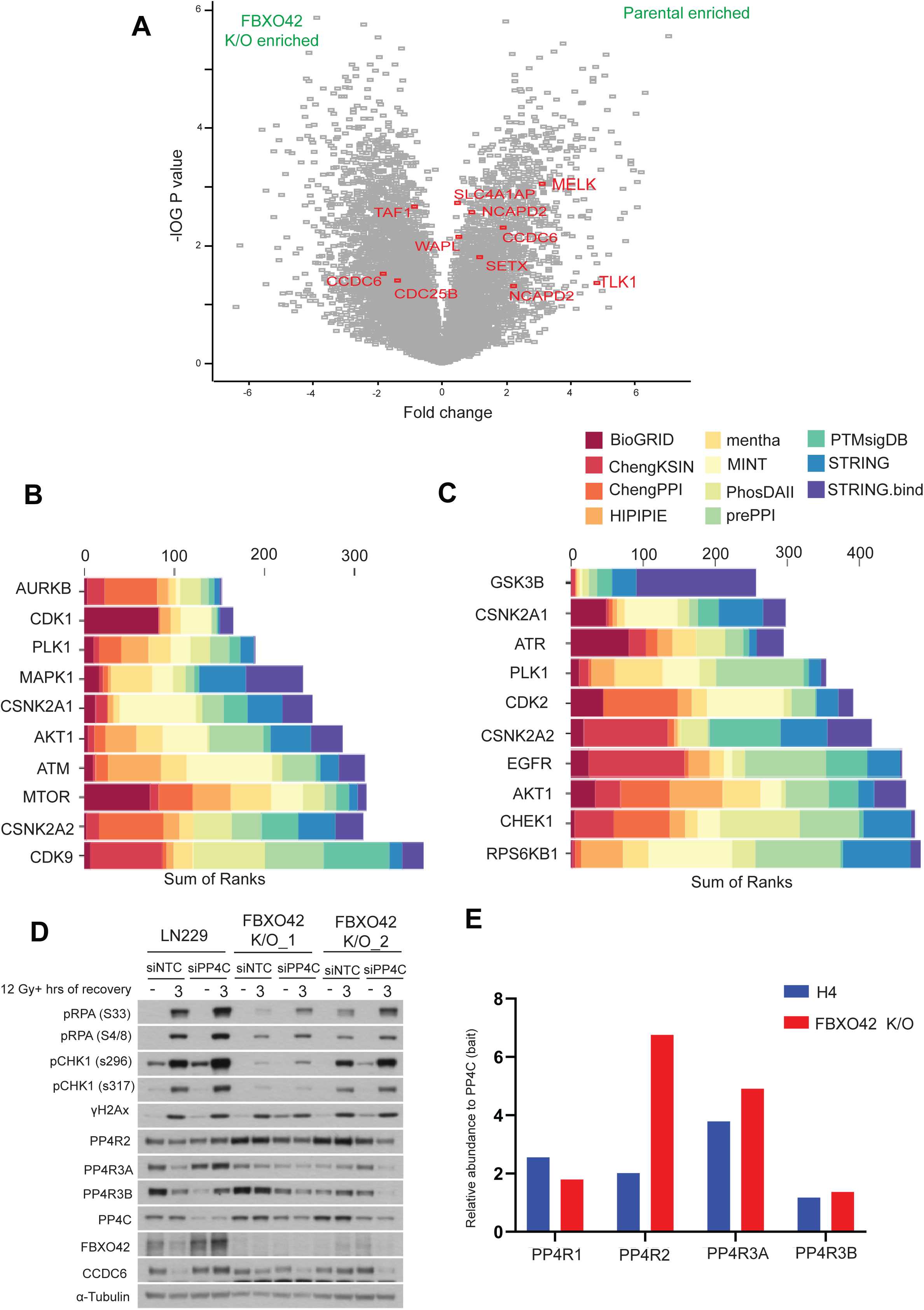
FBXO42 is a major regulator of PP4 phosphatase. A. Volcano plot depicting differentially enriched phosphorylated motifs / phosphosites in (total proteomes of) LN229 parental cells compared to Fbxo42 K/O cells. Consensus phosphosites in substrates of the PP4R2 subcomplex previously annotated by (Ueki et al., 2019) have been highlighted if considered significant using FDR=0.05 and S1=0.1 according to the Perseus software. B. Kinase enrichment analysis using KEA3 of proteins whose phosphorylation sites decreased in FBXO42 K/O cells when compared to LN229 parental. C. Kinase enrichment analysis using KEA3 of proteins whose phosphorylation sites increased in cells treated with siRNA against PP4C when compared to LN229 treated with siRNA control. D. Immunoblotting of DNA damage markers and PP4 complex subunits in LN299 parental and two independent FBXO42 K/O cell lines after IR treatment and 3-hour recovery with or without siRNA knockdown of PP4C. DNA damage markers detected: pRPA32 S33 (single-strand breaks), pRPA32 S4/8 (ssDNA at DSB), pCHK1 S296 (cell cycle checkpoint), pCHK1 S317 and γH2Ax (DNA DSBs). E. Relative abundance of PP4R1, PP4R2, PP4R3A and PP4R3B to PP4C (bait) determined by comparing LFQ Intensity (which is a measure of protein abundance) in H4 parental cell lines compared to H4 FBXO42 K/O after immunoprecipitation of Flag PP4C and mass spectrometry.

The activity of kinases is much better studied given the availability of specific chemical inhibitors and several studies to define their consensus phosphorylation sites (Cohen et al., 2021; Johnson et al., 2023). In our model loss of FBXO42 leads to activation of PP4 phosphatase and, thus, decreased phosphorylation. Thus, we queried the kinases whose activity decreased in FBXO42 K/O cells by using Kinase Enrichment Analysis 3 (KEA3), an online tool to infer the activity of kinases from a list of proteins. The kinases whose activity was reduced (counteracted by PP4) were AURKB, CDK1, PLK1, MAPK1, CSNK2A1, AKT1, ATM, MTOR, CSNK2A2 and CDK9 (Figure 4B). The reduction in kinase activity is in line with the phenotypes associated to FBXO42 depletion and these kinases phosphorylate known PP4 substrates (Park & Lee, 2020).

To establish a further association with the activity of PP4 we compared the kinase activity profiles from FBXO42 K/O cells to cells in which PP4 activity is curbed by siRNA, given the lack of specific inhibitors. In this case we input a list of proteins whose phosphorylation was increased in PP4C siRNA, into KEA3.

The kinases thus identified were GSK3B, CSNK2A1, ATR, PLK1, CDK2, CSNK2A2, EGFR, AKT1, CHEK1 and RPS6KB1 (Figure 4C). Out of the 10 kinases identified in FBXO42 K/O or PP4C siRNA, five were identical and the others had very closely related profiles (Figure 4B and 4C), again a strong indication that PP4 activity is altered in these cells. The lack of a stronger or perhaps even a complete overlap could be attributed to technical limitations including only a partial depletion of PP4C, the presence of indirect effects, and/or the presence of additional substrates modulated by the PP4R1 and PP4R4 complexes which are also impacted by PP4C depletion.

To consolidate multiple observations thus far, we aimed to understand whether phosphorylation events occurring upon IR treatment in FBXO42 lacking cells could be rescued by concomitant depletion of PP4C. FBXO42 K/O cells showed a significant decrease of RPA phosphorylation (Figure 1F and 1G) which are known to be controlled by PP4 activity (Lee et al., 2010). Most importantly, the phosphorylation statuses of RPA and Chk1 was rescued by depletion of PP4C in FBXO42 K/O (Figure 4D). This further consolidates the model whereby aberrant PP4 activity is brought about by the loss of FBXO42. Interestingly, the depletion of PP4R1 did not impact these phosphorylation events as much as the depletion of PP4R2, possibly indicating that the activity of FBXO42 after IR was mainly restricted to the PP4R2 subcomplexes (Figure S5A). A possible explanation for this observation is that the ubiquitination events on PP4C prevent binding to the PP4R2/R3A and PP4R2/R3B complexes. To confirm this, we tested for the ability of PP4C to bind PP4R2 in FBXO42 knockout cells. For this purpose, we immunoprecipitated PP4C from parental and FBXO42 K/O cells and measured the relative abundance of the PP4 associated subunit using mass spectrometry. We observed that the lack of FBXO42 increased the interaction of PP4R2 with PP4C (Figure 4E) but did not have the same impact on the interaction of PP4C with PP4R1. We also immunoprecipitated PP4R3A and PP4R3B in parental and FBXO42 K/O cells confirming an increase in the interaction observed with PP4C and PP4R2 (Figure S5B) – these subcomplexes being the true effectors of the observed phenotypes in FBXO42 knockout.

## Discussion

Here we identify an uncharacterised regulator of PP4 providing insights into an unprecedented regulatory mechanism. Our work indicates that the main function of FBXO42 is to restrict the activity of the PP4 phosphatase, particularly the PP4R2 complex. The mechanism is attributed to a likely non-degradative ubiquitination event operated by FBXO42 on PP4C to restrict the formation of active PP4C/PP4R2 complexes. Lack of FBXO42 unleashes uncontrolled PP4 activity thus incapacitating the action of kinases involved in cell cycle control and signalling, as shown by our phosphoproteomic approach. We observe that increased activity of PP4 in FBXO42 K/O results in defective DNA damage checkpoint signalling and reduced Chk1 and RPA phosphorylation. These events can be rescued by depletion of PP4 in an FBXO42 K/O background establishing a direct relationship between FBXO42 and PP4 in modulating DNA damage signalling (Figure 5). A manuscript with parallel observations points out that the survival of cancer cells depleted of FBXO42 can be reverted by concomitant depletion of PP4, solidifying the notion that FBXO42 act as a major regulator of PP4 (Spangenberg et al., 2025).

**Figure 5.**
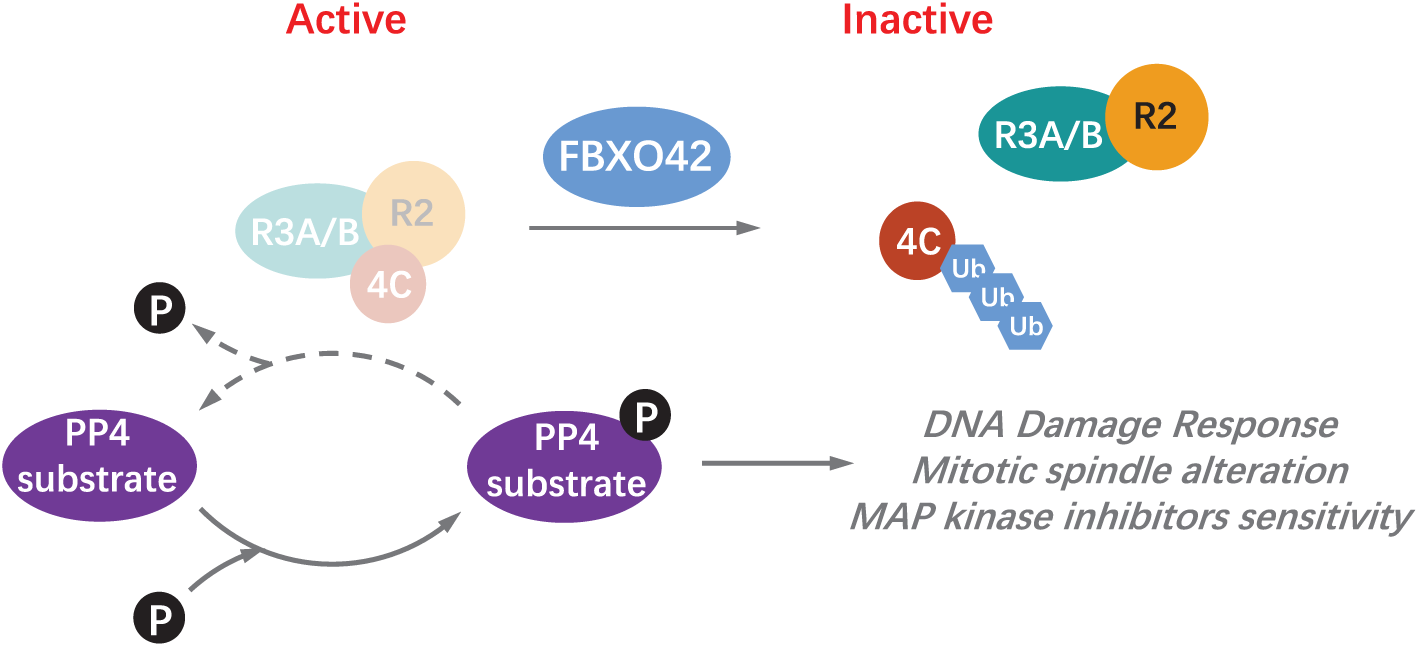
FBXO42 inhibits PP4 activity through ubiquitination of PP4C. **A.** Schematic depiction of the regulatory activity of FBXO42 on the PP4 complex.

The type of ubiquitination assembled by FBXO42 (on PP4) is likely to be non-degradative, but it is not possible to exclude that smaller pools of PP4C and/or PP4R2 are degraded in cells, thereby altering the stoichiometry of the PP4 subunits with the net effect of increasing PP4 activity. In the case of PP2A, at least five regulatory proteins are in place to assist the biogenesis of the full complex and restrict the activity of the catalytic subunit (alpha4, PTPA, LCMT1, PME-1 and TIPRL) (Fellner et al., 2003; Guo et al., 2014; Sents et al., 2013; Stanevich et al., 2011; Wu et al., 2017; Xing et al., 2008). FBXO42 might assist in the biogenesis of PP4 complexes to restrict activity until the full complex is assembled. Indeed, while classically C-terminal methylation of the catalytic subunit is required to assemble the PPP complexes, this is less important in the assembly of PP4C/PP4R2 where FBXO42 might play a major role (Hwang et al., 2016; Scott P. Lyons et al., 2021). At the moment, we can also not exclude that other PPP complexes are controlled by PP4, as we retrieve other phosphatases in our proteomic approach.

We provide initial evidence in FBXO42 K/O that the assembly of PP4C to the regulatory subunits is defective, however, further studies are required to establish the type of polyubiquitin chains formed by FBXO42 and how these chains mechanistically help the formation of PP4C/PP4R2 complexes. A recent study suggests RBPJ as a substrate of FBXO42, regulated through non degradative K63 ubiquitination (Jiang et al., 2022). We did not identify RBPJ in our interaction studies, however, it is plausible that FBXO42 catalyses non-degradative ubiquitin chains. It is unclear how mechanistically F-boxes can trigger non canonical ubiquitination having a common E2 recruiter in Rbx1.It may even be conceivable that super complex assemblies with E3s triggering an initial ubiquitination event, and other E3s catalysing chain extension might be in play (Scott et al., 2016). The paucity of reagents to detect non canonical ubiquitination strongly limit the capacity of further elucidating the details of the regulatory events controlled by non-degradative polyubiquitin chains.

Our work outlines a central regulatory node to control survival with several cancer associated phenotypes requiring further studies. FBXO42 inhibition leads to dysregulated kinase activity and might exacerbate a latent liability of cancer cells.

---

**Figure S1.**
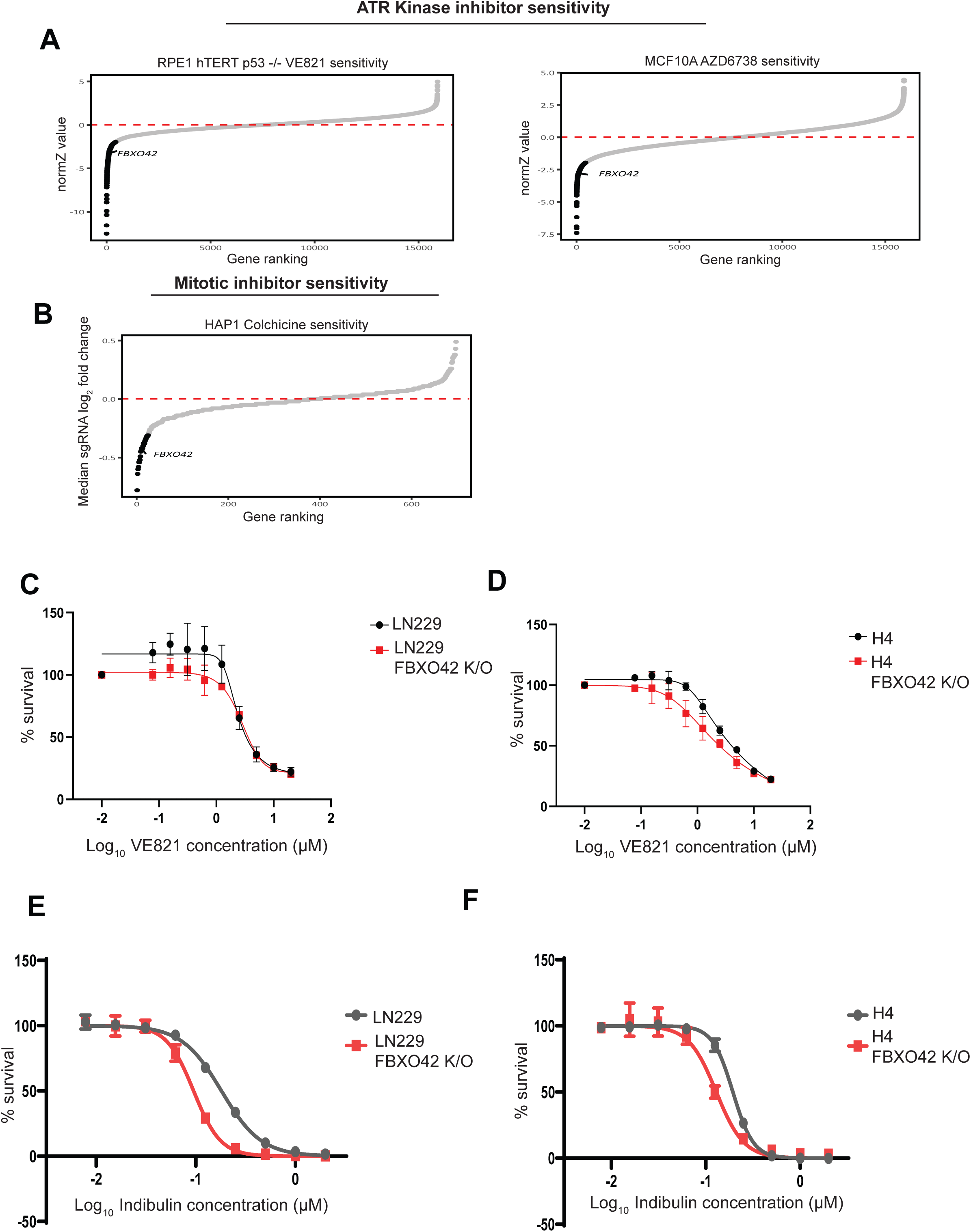
FBXO42 loss sensitises cells to ATR kinase inhibitors and microtubule depolimerising agents. A. CRISPR chemical-genetic screening results denoting vulnerabilities in response to ATR kinase inhibitors in RPE1 hTERT p53-/- and MCF10A cells (Hustedt et al., 2019; Wang et al., 2019), using a normZ significance cut-off of ≤ -1.96. FBXO42 is highlighted. B. CRISPR chemical-genetic screening results which details vulnerabilities in response to tubulin inhibitor colchicine in HAP1cells (Hundley et al., 2021), using a median sgRNA log2 fold change significance cut-off of ≤ -0.3. FBXO42 is highlighted. C. Dose response curves for ATR inhibitor VE821 to test the effect of FBXO42 K/O on cell survival in LN229 and H4 cell lines. D. Same as C in response to Indibulin.

**Figure S2.**
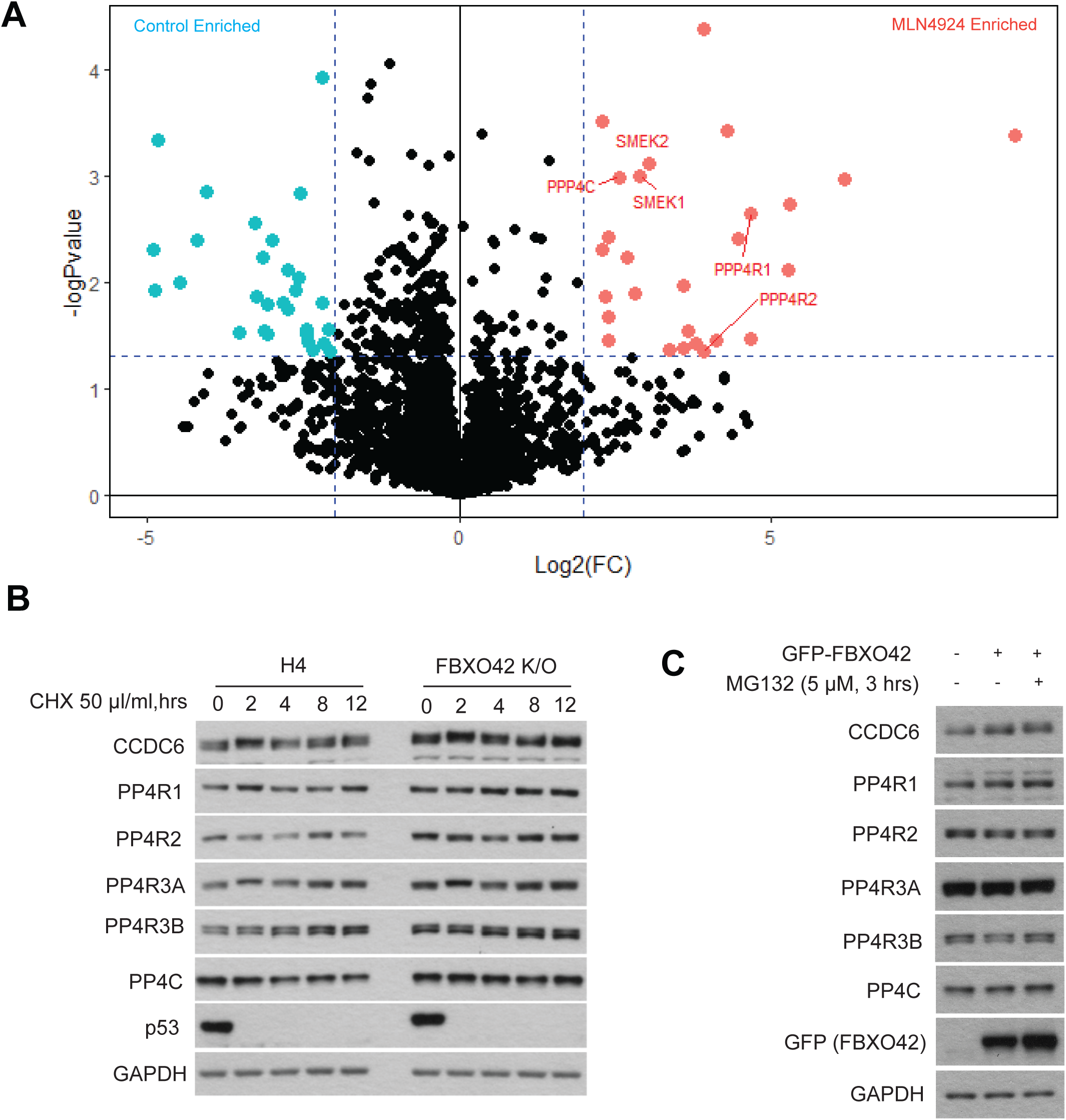
FBXO42 interacts with the PP4 complex. A. Volcano plot representing differentially enriched proteins (+/- MLN4924) generated using TurboID-FBXO42 followed by mass spectrometry. Interacting proteins were isolated with streptavidin after labelling for 1 hour with biotin. Proteins considered significant have a –log p value above 1.3 and log fold change of 2. In red are proteins enriched upon MLN4924, in blue enriched in untreated samples. B. Immunoblotting of FBXO42 interactors after treating H4 parental or FBXO42 K/O cells with cycloheximide (CHX) for the indicated times (hrs = hours). C. Immunoblotting of PP4 subunits after expression of GFP-FBXO42 in HEK293T with or without proteasomal inhibition with MG132 as indicated.

**Figure S3.**
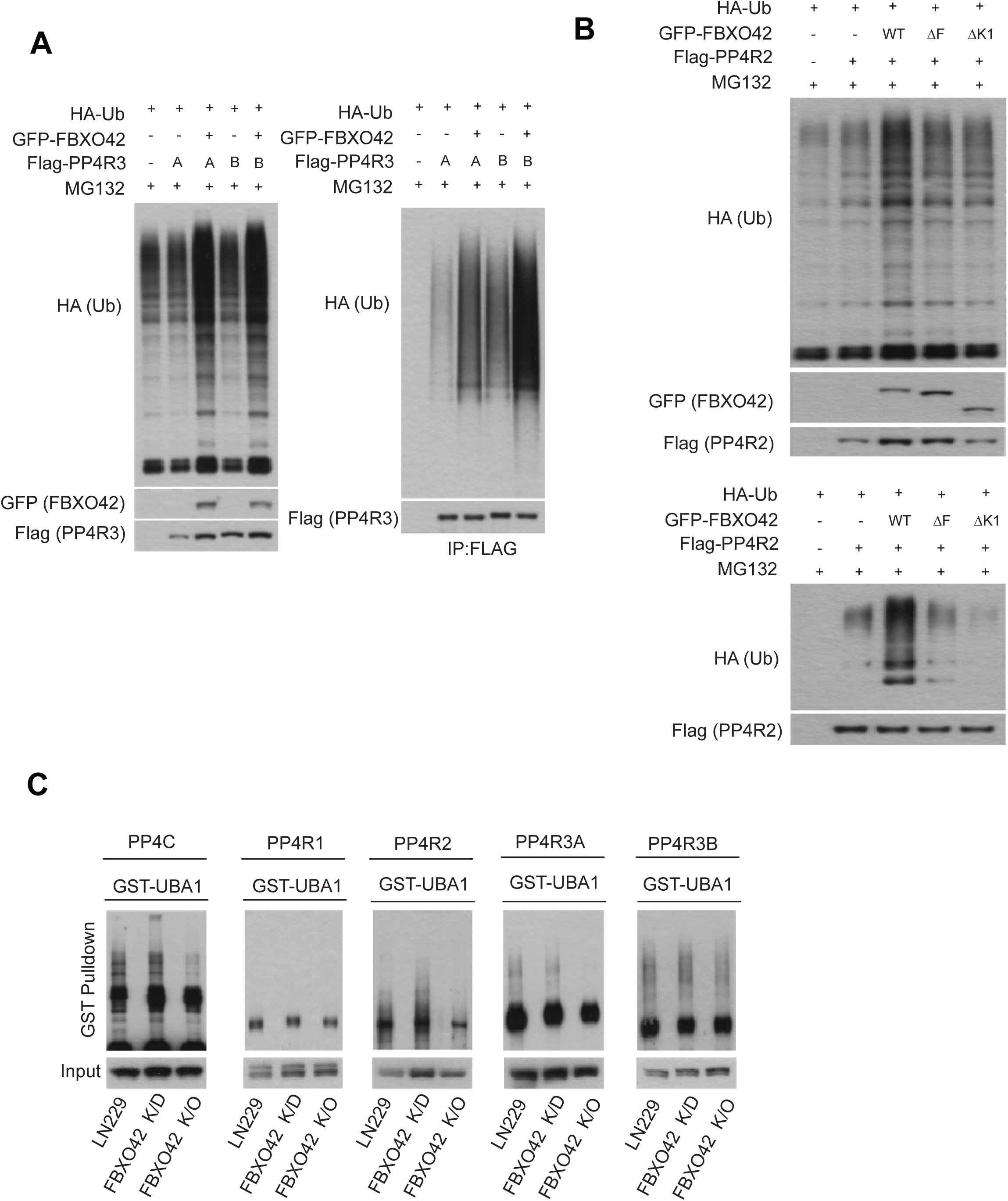
PP4 subunits are ubiquitinated by FBXO42. A. Immunoblotting after expression of GFP-FBXO42, HA-ubiquitin and either Flag-PP4R3A or Flag-PP4R3B in HEK293T. Flag-PP4R3A or Flag-PP4R3B (A or B) was isolated via Flag agarose beads pulldown before immunoblotting. Input samples are in the left panel. B. Immunoblotting after expression of HA-ubiquitin, Flag-PP4R2 and either GFP-FBXO42 WT, F-box (DF) mutant and Kelch (DK1) mutant in HEK293T. Flag-PP4R2 was isolated via Flag agarose beads pulldown before immunoblotting. Input samples are in the top panel. C. Immunoblotting after isolation of endogenous ubiquitinated proteins using recombinant GST-tagged UBA domain of UBIQLN protein from LN229 parental, FBXO42 K/D (partial knockout of FBXO42) or FBXO42 K/O cells. Input samples before immunoprecipitations are indicated in the bottom panel.

**Figure S4.**
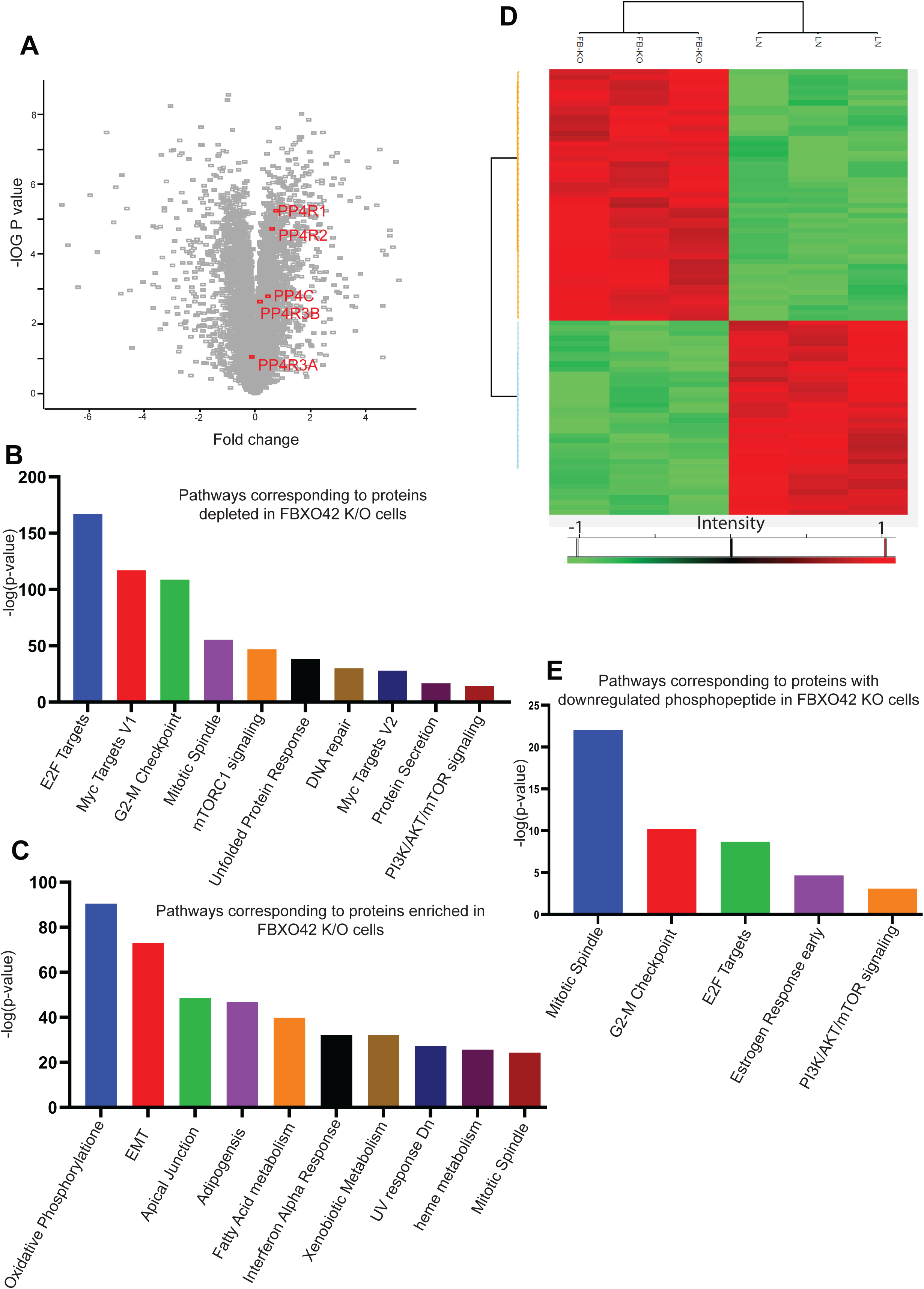
FBXO42 K/O mediated changes in total proteome and phosphoproteome. A. Volcano plot depicting differentially enriched phosphosites LN229 parental cells compared to FBXO42 K/O cells. Proteins containing consensus phosphosites in substrates of the PP4R2 subcomplex annotated by (Ueki et al., 2019) have been indicated. B. Gene enrichment analysis of proteins enriched in parental cells compared to FBXO42 K/O generated using the MSigDB hallmarks 2020 enrichment tool on EnrichR. C. Gene enrichment analysis of proteins depleted in LN229 parental cell lines compared to FBXO42 K/O generated using the MSigDB hallmarks 2020 enrichment tool on EnrichR. D. Heatmap comparing expression of phosphopeptides in LN229 parental and FBXO42 K/O cells. Colours represent intensity values with positive values in red and negative values in Green. E. Gene enrichment analysis of proteins who’s corresponding phosphosites were downregulated in LN229 FBXO42 K/O cell lines generated using the MSigDB hallmarks 2020 enrichment tool on EnrichR.

**Figure S5.**
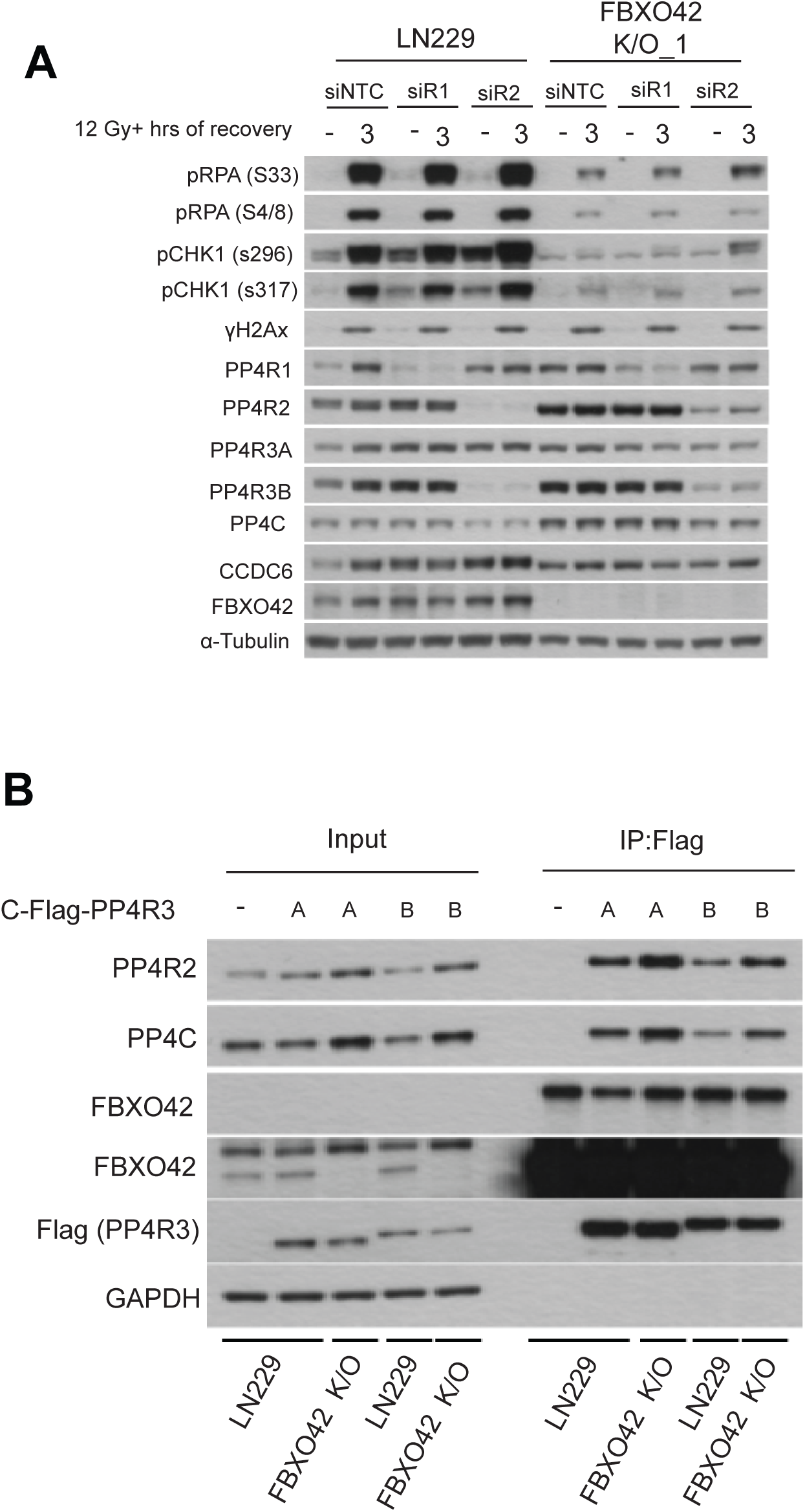
FBXO42 controls the activity of PP4 phosphatase and subunit assembly. A. Immunoblotting of DNA damage markers and PP4 complex subunits in LN299 parental and FBXO42 K/O cell lines after IR treatment and 3 hour recovery with or without siRNA knockdown of PP4R1 and PP4R2. DNA damage markers detected: pRPA32 S33 (single-strand breaks), pRPA32 S4/8 (ssDNA at DSB), pCHK1 S296 (cell cycle checkpoint), pCHK1 S317 and γH2Ax (DNA DSBs). B. Immunoblotting of endogenous PP4 subunits after expression of C-Flag-PP4R3A or Flag-PP4R3B in LN229 parental of FBXO42 K/O cell lines. Flag-PP4R3A/3B was isolated via Flag agarose beads pulldown before immunoblotting. Input samples are indicated.

## Author contributions

Conceptualization, investigation, methodology, resources, and experiment design: H.Y., V.D., P.S., B.M.K., R.F., I.V., D.G.; Ip/Wb experiments: H.Y., P.S., V.GK., B.S., J. R., Y.M.; phosphorylation study: H.Y., Y.M., P.S.; TurboID cell line and MS sample preparation/E-Stub: P.S.; bioinformatic analyses : E.S. ; writing and original draft: V.D., P.S., V.GK. ; proofreading, editing, and data curation: all authors; supervision: V.D., PS., H.Y., B.M.K., and R.F.

## Methods

### Reagents, antibodies and cell lines

All reagents, Antibodies and cell lines used in this study are listed in Table 1.

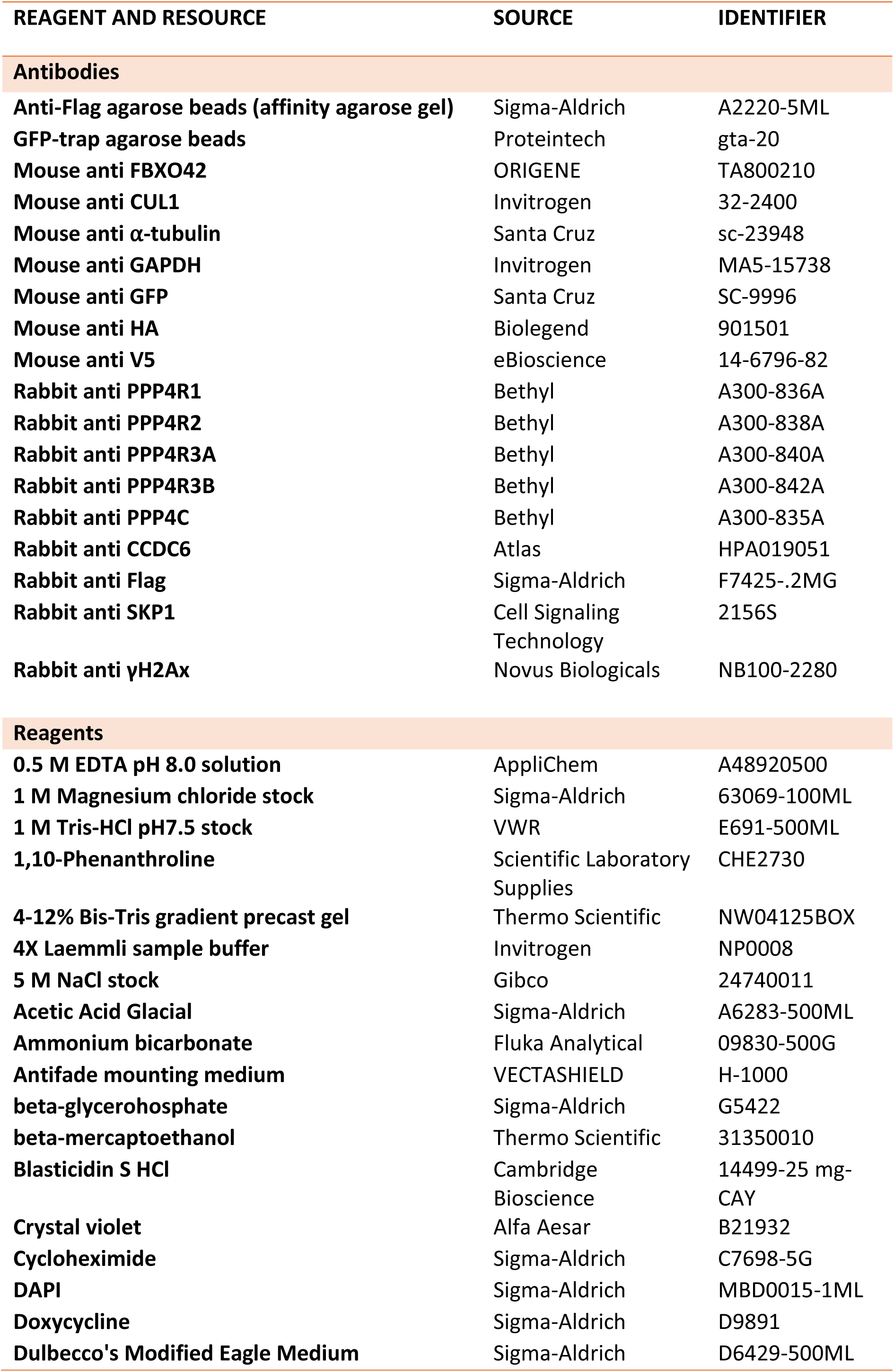
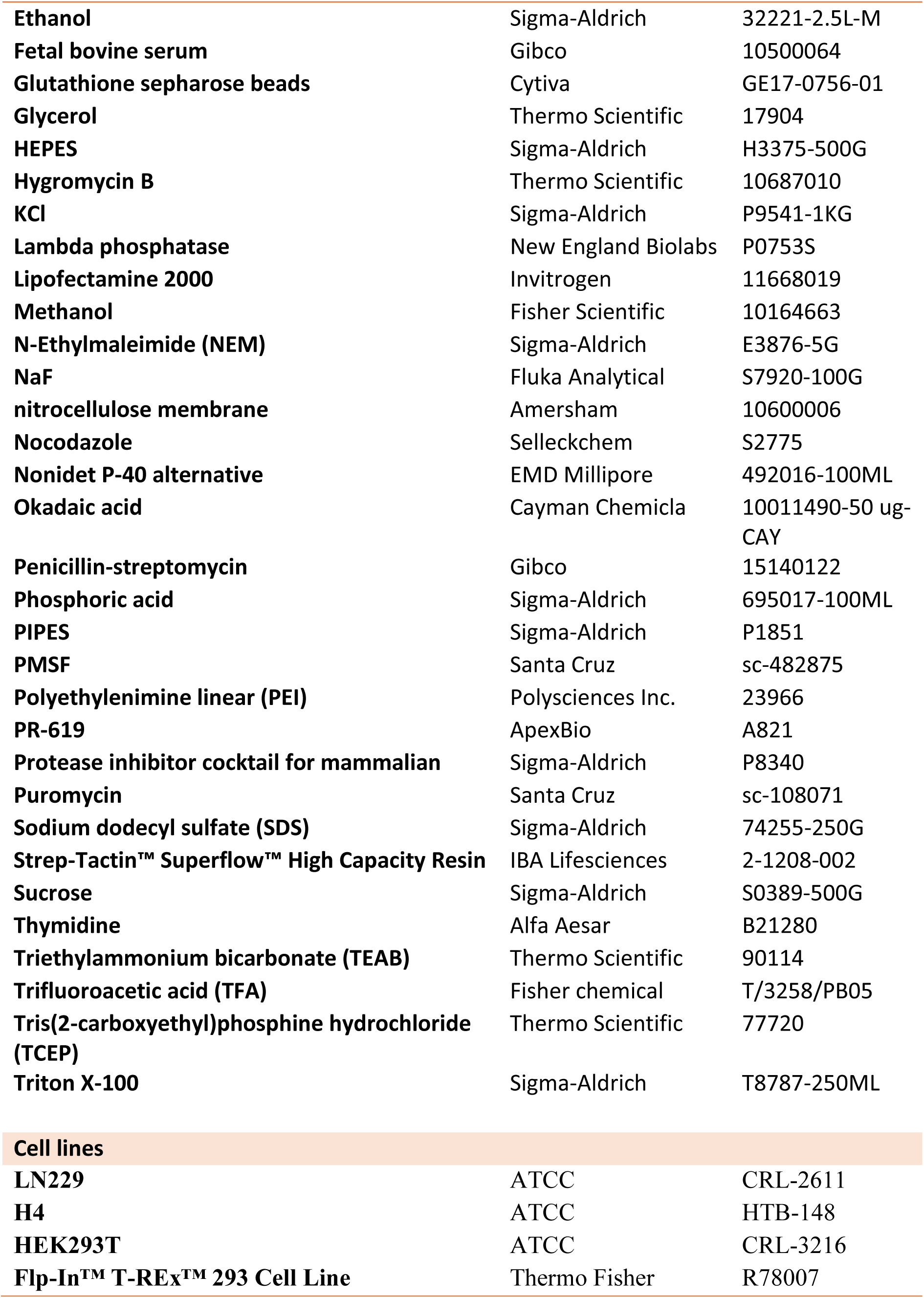

### Cell culture

LN229, HEK293T and H4 were obtained from American Type Culture Collection (ATCC). All cell lines were cultured in Dulbecco’s modified Eagle’s medium (DMEM; Sigma-Aldrich, D6429-500ML) containing 10% fetal bovine serum (FBS; Life Technologies, 10500064) and the mixture of penicillin (100 U/ml) and streptomycin (100 μg/ml; Life Technologies, 15140122). LN229 and H4 cells knockout for FBXO42 were generated using CRISPR (sgRNA sequences: table above).

### Colony formation assay

Cells were counted and seeded at 400 cells per well in six-well plates and cultured for 6 hours before being challenged with the indicated IR doses using a closed source gamma irradiator. After IR treatment, cells were allowed to propagate for 7 days (for HeLa-derived cell lines) or 14 days (for LN229-derived cell lines). Colonies were fixed and visualized using crystal violet solution (50% methanol, 10% ethanol, and 0.3% crystal violet) for 10 min at room temperature with gentle agitation (10 to 20 rpm). After rinsing with water and airdrying, colonies were counted using GelCount mammalian-cell colony counter (Oxford OPTRONIX). All colony formation assays presented in this study have been repeated at least three times and error bars represent SD of three biologically replicates. Statistical analysis was done using two-tailed unpaired t test.

### CRISPR knockout

Stable knockout was generated by co<u>-</u>transfecting Cas9 protein and three sgRNAs targeting the same gene, designed by Synthego, using Lipofectamine CRISPR Max (Life Technologies, CMAX00001) according to the protocol here: https://www.synthego.com/products/crispr-kits/synthetic-sgrna. Four days after transfection, cells were trypsinized and seeded as single cells in 96-well plates to isolate single clones. Cells in wells with proliferative clones were then trypsinized and expanded in bigger vessel until there were enough cells for immunoblotting. Clones that showed clear knockout were further validated by genomic DNA extraction and PCR amplification of the exon targeted by the sgRNAs. The PCR product was ligated into pCR4 vector using a TOPO TA cloning kit (Invitrogen, 450030). After transforming into DH5α competent cells, 10 colonies were picked and sent for Sanger sequencing. In LN229, cells were co<u>-</u> transfected with Cas9 protein and Synthego sgRNAs and directly subjected to treatment and western blotting (WB) 4 days after transfection. Although single clone picking was not performed, knockout efficiency in the mixed population was proven to be adequate.

### Generation of TurboID–FBXO42 stable cell line

Flp-In™ T-REx™ HEK293 cells (Invitrogen, R78007) containing a single genomic FRT site and stably expressing the Tet repressor were cultured in DMEM supplemented with 10% FBS, zeocin (100 μg/ml), and blasticidin (15 μg/ml).

The medium was exchanged with fresh medium containing no antibiotics before transfection. For cell line generation, Flp-In HEK293 cells were cotransfected with the pCDNA3–TurboID–FBXO42 plasmid and the pOG44 Flp– recombinase expression vector (Invitrogen, V600520) for coexpression of the Flp-recombinase using Lipofectamine 2000 transfection reagent (Invitrogen, 11668019). Two days after the transfection, cells were selected in hygromycin-containing medium (100 μg/ml) for 2 to 3 weeks. To validate the TurboID–FBXO42 expression, cells were cultured in media containing doxycycline (1.3 μg/ml) for 24 hours to induce TurboID–FBXO42 expression before immunoblotting.

### MS sample preparation for TurboID–FBXO42 pulldown

When TurboID–FBXO42 Flp-In™ T-REx™ HEK293 cells grown in 15-cm dishes reached 80% confluency, doxycycline (1.3 μg/ml) was added for 24 hours to induce the expression of TurboID–FBXO42. Cells were further incubated with 50 μM biotin for 3 hours to label proteins that came into close proximity with TurboID–FBXO42 in cells. Cells were harvested by scraping and washed three times with phosphate-buffered saline (PBS). For streptavidin pulldown of all biotin-labelled proteins (potential FBXO42 interactors), cell pellets were thoroughly resuspended in 1 ml of RIPA buffer [50 mM tris-HCl (pH 8.0), 150 mM NaCl, 1% Triton X-100, 1 mM EDTA, and 0.1% SDS with protease inhibitor cocktail (Sigma-Aldrich, P8340)] and incubated on ice for 15 min. Insoluble material was removed by centrifugation. Cleared lysates were then incubated on a rotating wheel at 4°C with 50-μl pre-equilibrated Strep-Tactin® Superflow® high capacity resin (IBA, 2-1208-002) for 1 hour. The suspension was then loaded on a Mini Bio-Spin Columns (Bio-Rad, 732-6207) to collect the beads. The beads were washed two times with 1 ml of RIPA buffer, three times with HNN buffer [50 mM HEPES (pH 7.5), 150 mM NaCl, and 50 mM NaF], and two times with 100 mM NH4HCO3 solution before being transferred to 2-ml Eppendorf tube in 400 μl of NH4HCO3 solution. For proteolysis, the sample was centrifuged at 200g for 1 min to remove supernatant. Beads were resuspended in 100 μl of 8 M Urea in 100 mM NH4HCO3 solution and incubated at 20°C for 20 min. Cysteine bonds were reduced with a final concentration of 5 mM tris(2-carboxyethyl) phosphine hydrochloride (TCEP) for 30 min at 37°C and alkylated in a final concentration of 10 mM iodoacetamide for 30 min at room temperature in the dark. Beads were then proteolyzed with trypsin/Lys-C Mix (Promega, V5071) at a 25:1 protein:protease ratio (w/w) for 4 hours at 37°C on an orbital shaker. Urea concentration was then reduced to 1 M via adding 100 mM NH4HCO3 solution to the sample. Samples were digested overnight at 37°C on an orbital shaker. Samples were desalted on C18 spin columns (Thermo Fisher Scientific, 89870) and washed according to the manual provided by the manufacturer and eluted with 0.1% trifluoroacetic acid (TFA) and 65% acetonitrile. Peptides were then dried in a SpeedVac vacuum concentrator and resuspend in 0.1% TFA and 2% acetonitrile in MS-grade water for MS analysis.

### Total proteome analysis

Heatmap of the total proteomes of LN229 and FBXO42 KO cells were analysed by hierarchical clustering using Euclidean distance and average for linkage method. The proteins from the clustering were used for pathway analysis using EnrichR.

### Phosphoproteomics analysis

Heatmap of the Phosphopeptides in LN229 and FBXO42 KO cells and siRNA PP4C were analysed by hierarchical clustering using Euclidean distance and average for linkage method. The proteins cluster upregulated in siPP4C and downregulated in FBXO42 KO cells were analysed using Kinase Enrichment Analysis version 3 (KSE3) to infer upstream kinases whose putative substrates are overrepresented in these differentially phosphorylated proteins. Pathway analysis was also done on the FBXO42 KO depleted phosphorylated proteins using EnrichR.

### Transfection, IP, and WB

A total of 1.5 million HEK293T cells were seeded into each of the 10-cm petri dish 24 hours before plasmid transfection. For each 10-cm dish, the polyethylenimine linear (PEI) transfection was performed by vigorously vortexing 5 μg of plasmid DNA with 15 μl of PEI (2.5 mg/ml) in 400 μl of plain DMEM (without FBS or antibiotics) for 15 s to mix, incubating the mixture for 15 min at room temperature, then adding it to cells cultured in 10-ml complete media in a dropwise fashion. PEI was purchased from Polysciences Inc. (23966). PEI stock (2.5 mg/ml) was made in 20 mM HEPES, 150 mM NaCl (pH 7.4), and filtered. Twenty-four hours after transfection, cells were washed twice with PBS and harvested. The cell pellets were stored at −80°C or lysed directly for experiments.

Cell pellets harvested for IP or WB not aiming to detect DNA damage markers were lysed in lysis buffer containing 50 mM tris-HCl (pH 7.5), 150 mM NaCl, 1 mM EDTA, 5 mM MgCl2, and 0.1% Nonidet P-40, supplemented with protease inhibitor cocktail (Sigma-Aldrich, P8340), 200 μM phenylmethylsulfonyl fluoride (PMSF; Santa Cruz Biotechnologies, sc-482875), and two phosphatase inhibitors, 20 mM β-glycerophosphate (Sigma-Aldrich, G5422) and 1 μM Okadaic acid (Cayman Chemical, 10011490-50 ug-CAY). After lysing on ice for 10 min, the insoluble fraction (mostly DNA and DNA bound proteins) was removed via centrifugation at 20,000 g at 4°C for 15 min, and the supernatant was carefully transferred to new Eppendorf tubes without disturbing the insoluble fraction. Protein concentration of the supernatant was measured using the modified Lowry assay (DC Protein Assay Kit, Bio-Rad, 5000111). The same amount of total protein was used for each IP (0.5 to 1 mg per pulldown in general) or direct WB (10 to 20 μg per lane in general). Final samples were mixed with 4X Laemmli sample buffer (Invitrogen, NP0008) and boiled for ten minutes before being applied in SDS-PAGE. For IP, 10-μl Flag M2 beads (Sigma-Aldrich, A2220-5ML) or 15-μl HA beads (Sigma-Aldrich, E6779-1ML) were washed three times with lysis buffer and added to the cell lysates to incubate for 3 hours on a roller at 4°C. After incubation, the beads were collected by centrifugation and washed with inhibitor-containing lysis buffer five times before being mixed with 20 <u>-</u>40 μl 1× Laemmli sample buffer (diluted from Invitrogen, NP0008) supplemented with β-mercaptoethanol, and boiled for 10 min at 95°C. The boiled supernatant can then be applied in WB analysis.

For experiments detecting DNA damage markers (which could be tightly chromatin bound), cell pellets were harvested, homogenized, and boiled directly in 2% SDS buffer [350 mM bis-tris (pH 6.8), 20% glycerol, and 2% SDS], then sonicated. Protein concentration was assessed using a BCA protein kit (Thermo Fisher Scientific, 23227). Cell lysate was prepared for WB as indicated above and resolved in 4 to 12% gradient bis-tris gels, transferred onto nitrocellulose membrane (Amersham, 10600006) and immunoblotted. WB results were visualized via x-ray film or iBright FL1500 Imaging System (Invitrogen, A44241).

### Cyclohexamide (CHX) chase

A total of 250,000 cells were seeded into each well of a six-well plate. Sixteen hours later, cells were cultured with CHX (50 μg/ml) for various durations as indicated in the figures, to block ribosomal protein synthesis for assessment of protein stability. Cells were collected via scraping and washed with PBS twice before being subjected immediately to WB for protein half-life estimation.

### In vivo ubiquitination assay

Two million HEK293T cells were seeded into each of the 10-cm dishes 24 hours before being transfected with plasmids as indicated in each experiment (typically, for each 10-cm dish, 1 μg of substrate overexpression plasmid, 2 μg of E3 overexpression plasmid, or 3 μg of ubiquitin overexpression plasmid was co<u>-</u>transfected). Cells were cultured for another 24 hours before being harvested. Four hours before the harvest, MG132 was added to a final concentration of 10 μM to block proteasomal degradation so that ubiquitination events were enriched. Cells were collected by scrapping and washed with PBS once, then thoroughly lysed and boiled in 300 μl of ubiquitin lysis buffer [2% SDS, 150 mM NaCl, and 10 mM tris-HCl (pH 7.4)]. After cooling to room temperature, cell lysates were subjected to sonication until they lost their viscosity. Lysates were then boiled again and centrifuged at 17,000 g for 10 min. Twenty μl of supernatant was preserved as input for each sample. The rest of the supernatant was diluted 20 times with dilution buffer [10 mM tris-HCl (pH 7.4), 150 mM NaCl, 2 mM EDTA, and 1% Triton X-100] and processed by IP of the substrate. After 16 hours of incubation on a roller at 4°C, beads were collected via centrifugation at 2000 rpm for 1 min and washed five times with 1 ml wash buffer [10 mM tris-HCl (pH 7.4), 1 M NaCl, 1 mM EDTA, and 1% Nonidet P-40] before being mixed with 50 μl of 1X Laemmli sample buffer, boiled, and subjected to WB.

### Ubiquitin binding entities (UBE) pulldown assay

Before harvesting cells, GST-UBA [ubiquitin-associated domain (UBA domain) of the UBQLN1 protein] was first conjugated to glutathione sepharose beads (Cytiva, GE17-0756-01) in ubiquitin binding entities (UBE) lysis buffer [19 mM NaH2PO4, 81 mM Na2HPO4 (pH 7.4), 1% Nonidet P-40, 2 mM EDTA, supplemented with protease inhibitor cocktail, PMSF, and phosphatase inhibitors as mentioned in the IP section], 50 mM N-ethylmaleimide (NEM; Sigma-Aldrich, E3876-5G), 5 mM 1,10-phenanthroline (Scientific Laboratory Supplies, CHE2730), and 50 μM PR-619 (ApexBio, A821) for at least 4 hours at 4°C on a roller. For each pulldown, 100 μg of recombinant GST-UBA was conjugated to 20 μl of washed glutathione beads. Cells were treated with 10 μM MG132 for 4 hours before being harvested by scrapping and centrifugation. Cell pellets were washed twice with PBS, then directly lysed in freshly prepared UBE lysis buffer. After incubation on ice for 10 min, lysates were subjected to centrifugation at 14,000 rpm at 4°C for 15 min. After protein concentration was measured via Lowry assay using DC Protein assay kit (Bio-Rad, 5000111), the same amount of total protein was used for each pulldown (2 to 3 mg per pulldown). GST-UBA–conjugated beads were added to the lysate and incubated on a roller at 4°C for overnight, then collected, washed using UBE lysis buffer, mixed with 1× Laemmli sample buffer, and boiled in the same way as described in the IP section. The supernatant was then used for WB. Besides the protein of interest, total ubiquitin was probed as a loading control for UBE pulldown experiments.

### Analysis of previous published CRISPR screens

Publicly available CRISPR-Cas9 screening results were extracted from publications concerning sensitivity to ataxia telangiectasia mutated and RAD3 related (ATR) protein kinase inhibitors AZD6738 and VE821 (Hustedt et al., 2019; Wang et al., 2019) (Panel A); mitotic inhibitors BI-2536 and colchicine (Hundley et al, 2021) (Panel B); and X-ray irradiation (Yang et al, 2024) (Panel C). Genes were ranked according to lowest normalized gene-level Z-score (normZ) value for Panel A, in which a significance cut-off value was defined as ≤ -1.96, lowest single guide RNA (sgRNA) median log2 fold change (Log2FC) for Panel B, in which a significance cut-off value was defined as ≤ -0.3 as per publication methods, and lowest Z-ratio for Panel C in which the significance cut-off value was lowered to ≤ -1 to reflect the use of focused CRISPR-Cas9 screening in place of genome wide.

### Generation of bio^GEF^ UBnc cell lines and bio^GEF^ UBnc+BIRA FBXO42 cell lines

Lentivirus expression construct TRIPZ-bio^GEF^ UBnc puro (TRIPZ-bio^GEF^-UBnc-PURO was a gift from Rosa Barrio & James Sutherland Addgene plasmid # 208044) were packaged in HEK293T cells by transfect with psPAX2, pMD2.G and pTAT (pcDNA1-Tat was a gift from Akitsu Hotta Addgene plasmid # 138478). Transfection media was removed after 12-16 hours and replaced with fresh media. The lentivirus supernatants were collected after 48 hours and filtered through 0.45 µM syringe filter. The lentivirus particles were used to transduce Flp-In™ T-®®™ 293 cells and antibiotic selection was performed with puromycin. To generate the bio^GEF^ UBnc+BIRA FBXO42 cell line the bio^GEF^ UBnc trex cell line were transfected with the pCDNA3–BIRA–FBXO42 plasmid and the pOG44 Flp–recombinase expression vector (Invitrogen, V600520) for coexpression of the Flp-recombinase using Lipofectamine 2000 transfection reagent (Invitrogen, 11668019). Two days after the transfection, cells were selected in hygromycin-containing medium (100 μg/ml) for 2 to 3 weeks. To validate the BIRA–FBXO42 expression, cells were cultured in media containing doxycycline (1.3 μg/ml) for 24 hours to induce BIRA-FBXO42 expression before immunoblotting.

### BioE3/E-STUB

BIRA–FBXO42 *bio^GEF^UBnc* Flp-In™ T-REx™ HEK293 cells grown in DMEM supplemented with 10% Tet FBS (PAN Biotech, P30-3602). One 15-cm dishes were used per experimental condition. When 15cm reached 80% confluency, doxycycline (1.3 μg/ml) was added for 24 hours to induce the expression of BIRA–FBXO42 and bio^GEF^UBnc. Cells were supplemented with 50 μM biotin for 3 hours before collection pre-treatment with MG132 or MLN4924 was 2 hours before biotin was added. Cells were harvested by scraping and washed three times with phosphate-buffered saline (PBS). For streptavidin pulldown cell pellets were thoroughly resuspended in lysis buffer (8M Urea (in 1xPBS) 1% SDS,50 µM NEM and Protease inhibitor cocktail (Sigma-Aldrich, P8340)) and incubated on ice for 15 min and sonicated 3 cycles of 30 sec on 30sec off.

Insoluble material was removed by centrifugation. Cleared lysates were incubated on a rotating wheel at 4°C with 50-μl pre-equilibrated Strep-Tactin<u>®</u> Superflow<u>®</u> high capacity resin (IBA, 2-1208-002) over night. The beads were washed 3 times in lysis buffer and 3 times in RIPA buffer. Beads were then resuspended in 2x Laemmli Sample Buffer (Biorad#1610747) and boiled at 95 °C for 5 min. Samples were then analysed by western blot.

## References

Agrata, R., & Komander, D. (2025). Ubiquitin-A structural perspective. Mol Cell, 85(2), 323–346. 10.1016/j.molcel.2024.12.015

Barroso-Gomila, O., Merino-Cacho, L., Muratore, V., Perez, C., Taibi, V., Maspero, E., Azkargorta, M., Iloro, I., Trulsson, F., Vertegaal, A. C. O., Mayor, U., Elortza, F., Polo, S., Barrio, R., & Sutherland, J. D. (2023). BioE3 identifies specific substrates of ubiquitin E3 ligases. Nat Commun, 14(1), 7656. 10.1038/s41467-023-43326-8

Bi, Y., Cui, D., Xiong, X., & Zhao, Y. (2021). The characteristics and roles of beta-TrCP1/2 in carcinogenesis. FEBS J, 288(11), 3351–3374. 10.1111/febs.15585

Burdova, K., Yang, H., Faedda, R., Hume, S., Chauhan, J., Ebner, D., Kessler, B. M., Vendrell, I., Drewry, D. H., Wells, C. I., Hatch, S. B., Dianov, G. L., Buffa, F. M., & D’Angiolella, V. (2019). E2F1 proteolysis via SCF-cyclin F underlies synthetic lethality between cyclin F loss and Chk1 inhibition. EMBO J, 38(20), e101443. 10.15252/embj.2018101443

Busino, L., Donzelli, M., Chiesa, M., Guardavaccaro, D., Ganoth, D., Dorrello, N. V., Hershko, A., Pagano, M., & Draetta, G. F. (2003). Degradation of Cdc25A by beta-TrCP during S phase and in response to DNA damage. Nature, 426(6962), 87–91. 10.1038/nature02082

Cardozo, T., & Pagano, M. (2004). The SCF ubiquitin ligase: insights into a molecular machine. Nat Rev Mol Cell Biol, 5(9), 739–751. 10.1038/nrm1471

Cerrato, A., Merolla, F., Morra, F., & Celetti, A. (2018). CCDC6: the identity of a protein known to be partner in fusion. Int J Cancer, 142(7), 1300–1308. 10.1002/ijc.31106

Chen, G. I., Tisayakorn, S., Jorgensen, C., D’Ambrosio, L. M., Goudreault, M., & Gingras, A. C. (2008). PP4R4/KIAA1622 forms a novel stable cytosolic complex with phosphoprotein phosphatase 4. J Biol Chem, 283(43), 29273–29284. 10.1074/jbc.M803443200

Chen, Z., Ioris, R. M., Richardson, S., Van Ess, A. N., Vendrell, I., Kessler, B. M., Buffa, F. M., Busino, L., Clifford, S. C., Bullock, A. N., & D’Angiolella, V. (2022). Disease-associated KBTBD4 mutations in medulloblastoma elicit neomorphic ubiquitylation activity to promote CoREST degradation. Cell Death Differ, 29(10), 1955–1969. 10.1038/s41418-022-00983-4

Cohen, P., Cross, D., & Jänne, P. A. (2021). Kinase drug discovery 20 years after imatinib: progress and future directions. Nature Reviews Drug Discovery, 20(7), 551–569. 10.1038/s41573-021-00195-4

Damgaard, R. B. (2021). The ubiquitin system: from cell signalling to disease biology and new therapeutic opportunities. Cell Death Differ, 28(2), 423–426. 10.1038/s41418-020-00703-w

Duda, D. M., Borg, L. A., Scott, D. C., Hunt, H. W., Hammel, M., & Schulman, B. A. (2008). Structural insights into NEDD8 activation of cullin-RING ligases: conformational control of conjugation. Cell, 134(6), 995–1006. 10.1016/j.cell.2008.07.022

Fellner, T., Lackner, D. H., Hombauer, H., Piribauer, P., Mudrak, I., Zaragoza, K., Juno, C., & Ogris, E. (2003). A novel and essential mechanism determining specificity and activity of protein phosphatase 2A (PP2A) in vivo. Genes Dev, 17(17), 2138–2150. 10.1101/gad.259903

Fiil, B. K., Damgaard, R. B., Wagner, S. A., Keusekotten, K., Fritsch, M., Bekker-Jensen, S., Mailand, N., Choudhary, C., Komander, D., & Gyrd-Hansen, M. (2013). OTULIN restricts Met1-linked ubiquitination to control innate immune signaling. Mol Cell, 50(6), 818–830. 10.1016/j.molcel.2013.06.004

Fouad, S., Wells, O. S., Hill, M. A., & D’Angiolella, V. (2019). Cullin Ring Ubiquitin Ligases (CRLs) in Cancer: Responses to Ionizing Radiation (IR) Treatment. Front Physiol, 10, 1144. 10.3389/fphys.2019.01144

Fung, E., Richter, C., Yang, H. B., Schaffer, I., Fischer, R., Kessler, B. M., Bassermann, F., & D’Angiolella, V. (2018). FBXL13 directs the proteolysis of CEP192 to regulate centrosome homeostasis and cell migration. EMBO Rep, 19(3). 10.15252/embr.201744799

Guardavaccaro, D., Kudo, Y., Boulaire, J., Barchi, M., Busino, L., Donzelli, M., Margottin-Goguet, F., Jackson, P. K., Yamasaki, L., & Pagano, M. (2003). Control of meiotic and mitotic progression by the F box protein beta-Trcp1 in vivo. Dev Cell, 4(6), 799–812. 10.1016/s1534-5807(03)00154-0

Guo, F., Stanevich, V., Wlodarchak, N., Sengupta, R., Jiang, L., Satyshur, K. A., & Xing, Y. (2014). Structural basis of PP2A activation by PTPA, an ATP-dependent activation chaperone. Cell Res, 24(2), 190–203. 10.1038/cr.2013.138

Harper, J. W., & Schulman, B. A. (2021). Cullin-RING Ubiquitin Ligase Regulatory Circuits: A Quarter Century Beyond the F-Box Hypothesis. Annu Rev Biochem, 90, 403–429. 10.1146/annurev-biochem-090120-013613

Hershko, A., & Ciechanover, A. (1992). The ubiquitin system for protein degradation. Annu Rev Biochem, 61, 761–807. 10.1146/annurev.bi.61.070192.003553

Hoellerbauer, P., Kufeld, M., Arora, S., Mitchell, K., Girard, E. J., Herman, J. A., Olson, J. M., & Paddison, P. J. (2024). FBXO42 activity is required to prevent mitotic arrest, spindle assembly checkpoint activation and lethality in glioblastoma and other cancers. NAR Cancer, 6(2), zcae021. 10.1093/narcan/zcae021

Huang, H.-T., Lumpkin, R. J., Tsai, R. W., Su, S., Zhao, X., Xiong, Y., Chen, J., Mageed, N., Donovan, K. A., Fischer, E. S., & Sellers, W. R. (2024). Ubiquitin-specific proximity labeling for the identification of E3 ligase substrates. Nature Chemical Biology, 20(9), 1227–1236. 10.1038/s41589-024-01590-9

Hundley, F. V., Sanvisens Delgado, N., Marin, H. C., Carr, K. L., Tian, R., & Toczyski, D. P. (2021). A comprehensive phenotypic CRISPR-Cas9 screen of the ubiquitin pathway uncovers roles of ubiquitin ligases in mitosis. Mol Cell, 81(6), 1319–1336 e1319. 10.1016/j.molcel.2021.01.014

Hustedt, N., Alvarez-Quilon, A., McEwan, A., Yuan, J. Y., Cho, T., Koob, L., Hart, T., & Durocher, D. (2019). A consensus set of genetic vulnerabilities to ATR inhibition. Open Biol, 9(9), 190156. 10.1098/rsob.190156

Hwang, J., Lee, J. A., & Pallas, D. C. (2016). Leucine Carboxyl Methyltransferase 1 (LCMT-1) Methylates Protein Phosphatase 4 (PP4) and Protein Phosphatase 6 (PP6) and Differentially Regulates the Stable Formation of Different PP4 Holoenzymes. J Biol Chem, 291(40), 21008–21019. 10.1074/jbc.M116.739920

Jiang, H., Bian, W., Sui, Y., Li, H., Zhao, H., Wang, W., & Li, X. (2022). FBXO42 facilitates Notch signaling activation and global chromatin relaxation by promoting K63-linked polyubiquitination of RBPJ. Sci Adv, 8(38), eabq4831. 10.1126/sciadv.abq4831

Jin, J., Shirogane, T., Xu, L., Nalepa, G., Qin, J., Elledge, S. J., & Harper, J. W. (2003). SCFbeta-TRCP links Chk1 signaling to degradation of the Cdc25A protein phosphatase. Genes Dev, 17(24), 3062–3074. 10.1101/gad.1157503

Johnson, J. L., Yaron, T. M., Huntsman, E. M., Kerelsky, A., Song, J., Regev, A., Lin, T.-Y., Liberatore, K., Cizin, D. M., Cohen, B. M., Vasan, N., Ma, Y., Krismer, K., Robles, J. T., van de Kooij, B., van Vlimmeren, A. E., Andrée-Busch, N., Käufer, N. F., Dorovkov, M. V., . . . Cantley, L. C. (2023). An atlas of substrate specificities for the human serine/threonine kinome. Nature, 613(7945), 759–766. 10.1038/s41586-022-05575-3

Komander, D., & Rape, M. (2012). The ubiquitin code. Annu Rev Biochem, 81, 203–229. 10.1146/annurev-biochem-060310-170328

Larochelle, M., Bergeron, D., Arcand, B., & Bachand, F. (2019). Proximity-dependent biotinylation mediated by TurboID to identify protein-protein interaction networks in yeast. J Cell Sci, 132(11). 10.1242/jcs.232249

Lee, D.-H., Pan, Y., Kanner, S., Sung, P., Borowiec, J. A., & Chowdhury, D. (2010). A PP4 phosphatase complex dephosphorylates RPA2 to facilitate DNA repair via homologous recombination. Nature Structural & Molecular Biology, 17(3), 365–372. 10.1038/nsmb.1769

Lu, Y., Cho, T., Mukherjee, S., Suarez, C. F., Gonzalez-Foutel, N. S., Malik, A., Martinez, S., Dervovic, D., Oh, R. H., Langille, E., Al-Zahrani, K. N., Hoeg, L., Lin, Z. Y., Tsai, R., Mbamalu, G., Rotter, V., Ashton-Prolla, P., Moffat, J., Chemes, L. B., . . . Schramek, D. (2024). Genome-wide CRISPR screens identify novel regulators of wild-type and mutant p53 stability. Mol Syst Biol, 20(6), 719–740. 10.1038/s44320-024-00032-x

Lyons, S. P., Greiner, E. C., Cressey, L. E., Adamo, M. E., & Kettenbach, A. N. (2021). Regulation of PP2A, PP4, and PP6 holoenzyme assembly by carboxyl-terminal methylation. Scientific Reports, 11(1), 23031. 10.1038/s41598-021-02456-z

Lyons, S. P., Greiner, E. C., Cressey, L. E., Adamo, M. E., & Kettenbach, A. N. (2021). Regulation of PP2A, PP4, and PP6 holoenzyme assembly by carboxyl-terminal methylation. Sci Rep, 11(1), 23031. 10.1038/s41598-021-02456-z

Margottin-Goguet, F., Hsu, J. Y., Loktev, A., Hsieh, H. M., Reimann, J. D., & Jackson, P. K. (2003). Prophase destruction of Emi1 by the SCF(betaTrCP/Slimb) ubiquitin ligase activates the anaphase promoting complex to allow progression beyond prometaphase. Dev Cell, 4(6), 813–826. 10.1016/s1534-5807(03)00153-9

May, D. G., & Roux, K. J. (2019). BioID: A Method to Generate a History of Protein Associations. Methods Mol Biol, 2008, 83–95. 10.1007/978-1-4939-9537-0_7

Merino-Cacho, L., Barroso-Gomila, O., Pozo-Rodriguez, M., Muratore, V., Guinea-Perez, C., Serrano, A., Perez, C., Cano-Lopez, S., Urcullu, A., Azkargorta, M., Iloro, I., Galdeano, C., Juarez-Jimenez, J., Mayor, U., Elortza, F., Barrio, R., & Sutherland, J. D. (2025). Cullin-RING ligase BioE3 reveals molecular-glue-induced neosubstrates and rewiring of the endogenous Cereblon ubiquitome. Cell Commun Signal, 23(1), 101. 10.1186/s12964-025-02091-5

Merino-Cacho, L., Barroso-Gomila, O., Pozo-Rodríguez, M., Muratore, V., Guinea-Pérez, C., Serrano, Á., Pérez, C., Cano-López, S., Urcullu, A., Azkargorta, M., Iloro, I., Galdeano, C., Juárez-Jiménez, J., Mayor, U., Elortza, F., Barrio, R., & Sutherland, J. D. (2025). Cullin-RING ligase BioE3 reveals molecular-glue-induced neosubstrates and rewiring of the endogenous Cereblon ubiquitome. Cell Communication and Signaling, 23(1), 101. 10.1186/s12964-025-02091-5

Morreale, F. E., & Walden, H. (2016). Types of Ubiquitin Ligases. Cell, 165(1), 248–248 e241. 10.1016/j.cell.2016.03.003

Nagler, A., Vredevoogd, D. W., Alon, M., Cheng, P. F., Trabish, S., Kalaora, S., Arafeh, R., Goldin, V., Levesque, M. P., Peeper, D. S., & Samuels, Y. (2020). A genome-wide CRISPR screen identifies FBXO42 involvement in resistance toward MEK inhibition in NRAS-mutant melanoma. Pigment Cell Melanoma Res, 33(2), 334–344. 10.1111/pcmr.12825

Ngoi, P., Wang, X., Putta, S., Da Luz, R. F., Serrao, V. H. B., Emanuele, M. J., & Rubin, S. M. (2025). Structural mechanism for recognition of E2F1 by the ubiquitin ligase adaptor Cyclin F. bioRxiv. 10.1101/2025.01.15.633208

Park, J., & Lee, D. H. (2020). Functional roles of protein phosphatase 4 in multiple aspects of cellular physiology: a friend and a foe. BMB Rep, 53(4), 181–190. 10.5483/BMBRep.2020.53.4.019

Raducu, M., Fung, E., Serres, S., Infante, P., Barberis, A., Fischer, R., Bristow, C., Thezenas, M. L., Finta, C., Christianson, J. C., Buffa, F. M., Kessler, B. M., Sibson, N. R., Di Marcotullio, L., Toftgard, R., & D’Angiolella, V. (2016). SCF (Fbxl17) ubiquitylation of Sufu regulates Hedgehog signaling and medulloblastoma development. EMBO J, 35(13), 1400–1416. 10.15252/embj.201593374

Saldivar, J. C., Cortez, D., & Cimprich, K. A. (2017). The essential kinase ATR: ensuring faithful duplication of a challenging genome. Nat Rev Mol Cell Biol, 18(10), 622–636. 10.1038/nrm.2017.67

Saldivar, J. C., Hamperl, S., Bocek, M. J., Chung, M., Bass, T. E., Cisneros-Soberanis, F., Samejima, K., Xie, L., Paulson, J. R., Earnshaw, W. C., Cortez, D., Meyer, T., & Cimprich, K. A. (2018). An intrinsic S/G(2) checkpoint enforced by ATR. Science, 361(6404), 806–810. 10.1126/science.aap9346

Scott, D. C., Rhee, D. Y., Duda, D. M., Kelsall, I. R., Olszewski, J. L., Paulo, J. A., de Jong, A., Ovaa, H., Alpi, A. F., Harper, J. W., & Schulman, B. A. (2016). Two Distinct Types of E3 Ligases Work in Unison to Regulate Substrate Ubiquitylation. Cell, 166(5), 1198–1214 e1124. 10.1016/j.cell.2016.07.027

Sents, W., Ivanova, E., Lambrecht, C., Haesen, D., & Janssens, V. (2013). The biogenesis of active protein phosphatase 2A holoenzymes: a tightly regulated process creating phosphatase specificity. FEBS J, 280(2), 644–661. 10.1111/j.1742-4658.2012.08579.x

Shi, Y. (2009). Serine/threonine phosphatases: mechanism through structure. Cell, 139(3), 468–484. 10.1016/j.cell.2009.10.006

Soucy, T. A., Smith, P. G., Milhollen, M. A., Berger, A. J., Gavin, J. M., Adhikari, S., Brownell, J. E., Burke, K. E., Cardin, D. P., Critchley, S., Cullis, C. A., Doucette, A., Garnsey, J. J., Gaulin, J. L., Gershman, R. E., Lublinsky, A. R., McDonald, A., Mizutani, H., Narayanan, U., . . . Langston, S. P. (2009). An inhibitor of NEDD8-activating enzyme as a new approach to treat cancer. Nature, 458(7239), 732–736. 10.1038/nature07884

Spangenberg, S. H., Garaffo, N., Catherine Alcindor, E. M., Lusk, B., Pandey, V., Longhurst, A., Bolech, E., Hua Fu, B. X., Gilbert, L. A., Wohlschlegel, J. A., & Toczyski, D. (2025). The regulation of Protein Phosphatase 4 by FBXO42 is required for cancer cell survival. bioRxiv, 2025.2004.2022.649889. 10.1101/2025.04.22.649889

Stanevich, V., Jiang, L., Satyshur, K. A., Li, Y., Jeffrey, P. D., Li, Z., Menden, P., Semmelhack, M. F., & Xing, Y. (2011). The structural basis for tight control of PP2A methylation and function by LCMT-1. Mol Cell, 41(3), 331–342. 10.1016/j.molcel.2010.12.030

Toledo, C. M., Ding, Y., Hoellerbauer, P., Davis, R. J., Basom, R., Girard, E. J., Lee, E., Corrin, P., Hart, T., Bolouri, H., Davison, J., Zhang, Q., Hardcastle, J., Aronow, B. J., Plaisier, C. L., Baliga, N. S., Moffat, J., Lin, Q., Li, X. N., . . . Paddison, P. J. (2015). Genome-wide CRISPR-Cas9 Screens Reveal Loss of Redundancy between PKMYT1 and WEE1 in Glioblastoma Stem-like Cells. Cell Rep, 13(11), 2425–2439. 10.1016/j.celrep.2015.11.021

Ueki, Y., Kruse, T., Weisser, M. B., Sundell, G. N., Larsen, M. S. Y., Mendez, B. L., Jenkins, N. P., Garvanska, D. H., Cressey, L., Zhang, G., Davey, N., Montoya, G., Ivarsson, Y., Kettenbach, A. N., & Nilsson, J. (2019). A Consensus Binding Motif for the PP4 Protein Phosphatase. Mol Cell, 76(6), 953–964 e956. 10.1016/j.molcel.2019.08.029

Wang, C., Wang, G., Feng, X., Shepherd, P., Zhang, J., Tang, M., Chen, Z., Srivastava, M., McLaughlin, M. E., Navone, N. M., Hart, G. T., & Chen, J. (2019). Genome-wide CRISPR screens reveal synthetic lethality of RNASEH2 deficiency and ATR inhibition. Oncogene, 38(14), 2451–2463. 10.1038/s41388-018-0606-4

Wu, C. G., Zheng, A., Jiang, L., Rowse, M., Stanevich, V., Chen, H., Li, Y., Satyshur, K. A., Johnson, B., Gu, T. J., Liu, Z., & Xing, Y. (2017). Methylation-regulated decommissioning of multimeric PP2A complexes. Nat Commun, 8(1), 2272. 10.1038/s41467-017-02405-3

Xing, Y., Li, Z., Chen, Y., Stock, J. B., Jeffrey, P. D., & Shi, Y. (2008). Structural mechanism of demethylation and inactivation of protein phosphatase 2A. Cell, 133(1), 154–163. 10.1016/j.cell.2008.02.041

Yang, H., Fouad, S., Smith, P., Bae, E. Y., Ji, Y., Lan, X., Van Ess, A., Buffa, F. M., Fischer, R., Vendrell, I., Kessler, B. M., & D’Angiolella, V. (2024). Cyclin F-EXO1 axis controls cell cycle-dependent execution of double-strand break repair. Sci Adv, 10(32), eado0636. 10.1126/sciadv.ado0636

Zheng, N., Schulman, B. A., Song, L., Miller, J. J., Jeffrey, P. D., Wang, P., Chu, C., Koepp, D. M., Elledge, S. J., Pagano, M., Conaway, R. C., Conaway, J. W., Harper, J. W., & Pavletich, N. P. (2002). Structure of the Cul1-Rbx1-Skp1-F boxSkp2 SCF ubiquitin ligase complex. Nature, 416(6882), 703–709. 10.1038/416703a

